# Nuclear DNA Damage is a Primary Driver of Mitochondrial Dysfunction in *C9ORF72* ALS/FTD

**DOI:** 10.64898/2026.05.20.726184

**Authors:** Mara Zilocchi, Illari Salvatori, Silvia Lombardi, Roland Nicsanu, Alice Campana, Roman Shaposhnikov, Gabriella Gualtieri, Silvia Scaricamazza, Cristiana Valle, Alberto Ferri, Silvia M.L. Barabino

## Abstract

Amyotrophic lateral sclerosis (ALS) is characterized by progressive motor neuron degeneration associated with genomic instability and mitochondrial dysfunction, although the mechanistic relationship between these hallmarks remains unclear.

To determine whether nuclear DNA damage alone induces mitochondrial dysfunction, we exploited the AID-DIvA system, selectively generating DNA double-strand breaks (DSBs) in nuclear DNA. DSB induction caused early impairment of mitochondrial bioenergetics, including reduced basal respiration, ATP-linked respiration, and maximal respiratory capacity, preceding more pronounced mitochondrial alterations, after prolonged damage. Resolution of DSBs restored mitochondrial function, demonstrating a direct and reversible link between nuclear genome instability and mitochondrial dysfunction. Mechanistically, persistent activation of the DNA damage response (DDR) triggered PARP1-dependent NAD□ depletion, while PARP1 inhibition rescued mitochondrial respiration and ATP synthesis.

We next investigated the consequences of DDR activation triggered by the expression of 102 (G4C2) repeats in an inducible cell model of C9ORF72-linked ALS. In these cells, DDR activation preceded mitochondrial dysfunction, recapitulating the sequence observed in AID-DIvA cells. Mitochondrial defects included impaired oxidative phosphorylation and reduced ATP production without increased mitochondrial ROS, suggesting that DNA damage signalling acts upstream of mitochondrial dysfunction. In support of this hypothesis, inhibition of ATM as well as nicotinamide riboside-mediated replenishment of cellular NAD^+^significantly restored mitochondrial functions.

Collectively, our findings identify nuclear DNA damage as a trigger of mitochondrial dysfunction and uncover a pathogenic DDR–mitochondria crosstalk mediated by persistent DNA damage signalling. These results support a bidirectional relationship between genome instability and mitochondrial dysfunction and highlight mitochondrial and DNA damage response modulators as potential therapeutic targets for ALS and related neurodegenerative disorders.

## Introduction

Amyotrophic lateral sclerosis (ALS) is an adult-onset progressive disorder caused by the selective death of motor neurons in the cortex, brainstem and spinal cord, leading to progressive muscle weakness, paralysis and ultimately death from respiratory failure. 10% of ALS cases are inherited and referred to as familial ALS (fALS). Currently more than 40 genes associated with fALS have been identified (Nijs & Van Damme, 2024). An intronic hexanucleotide G4C2 repeat expansion in C9ORF72 gene is the most common mutation in ALS and accounts for 5%–10% cases(Balendra & Isaacs, 2018).

To date a common molecular mechanism underlying neurodegeneration in ALS remains uncertain. Well-established hallmarks of ALS, contributing to vulnerability and degeneration of motor neurons, are genomic instability (Jagaraj et al., 2024), defective energy metabolism (Tefera & Borges, 2017), and mitochondrial dysfunction (MD)(Smith et al., 2019).

Even under basal energy demands, mitochondria are the main source of both ATP and reactive oxygen species (ROS), the majority of which are produced during oxidative phosphorylation. Once ROS are overproduced, they may lead to oxidative stress. The detection of increased oxidative DNA damage in affected tissues — often associated with defective mitochondrial metabolism (Bogdanov et al., 2000) — suggests that mitochondrial ROS-induced nuclear DNA damage may be central to ALS aetiology, placing MD upstream of genomic instability. However, recent evidence links many fALS-associated genes to the pathological cellular response induced by DNA damage (Konopka & Atkin, 2022; Sun et al., 2020), suggesting that DNA damage accumulation may be one of the main contributors to neurodegeneration in ALS.

Mammalian cells have evolved a complex cellular response, referred to as DNA Damage Response (DDR) to counteract DNA damage. The DDR coordinates DNA repair with signalling pathways regulating cell cycle progression and cell fate decisions. Upon DNA damage, sensors such as Poly(ADP-ribosyl) polymerase-1 (PARP-1) and PIKK kinases (DNA-PK, ATM, ATR) are rapidly activated leading to chromatin modifications. Importantly, DDR activation is tightly linked to cellular metabolic status and may impact mitochondrial function (Cucchi et al., 2021; Ma & Zhou, 2025).

Given the consolidated evidence that abnormal mitochondrial metabolism and defects in DNA damage repair are involved in neurodegenerative diseases, a still unresolved question is whether damage to nuclear DNA is sufficient to cause mitochondrial dysfunction. This question explicitly inverts the classical model in which mitochondrial dysfunction is considered the primary driver of nuclear DNA damage and instead tests a potential bidirectional relationship between nuclear genome instability and mitochondrial metabolism. However, the fact that several DNA repair-deficient premature ageing diseases, such as Cockayne syndrome, Xeroderma pigmentosum, and Ataxia telangiectasia, were shown to involve alterations in mitochondrial metabolism, suggesting that mitochondrial dysfunction may be a downstream consequence of the accumulation of nuclear DNA damage (Farg et al., 2017).

Here, we found that the specific induction of DNA double strand breaks (DSBs) in the genomic DNA is associated with significant alterations of mitochondrial respiratory parameters, increased parylation activity and depletion of cellular NAD+. These observations support the hypothesis that nuclear DNA damage may be sufficient to initiate MD even in the absence of direct mitochondrial genotoxic stress.

In addition, exploiting an inducible cell model of C9ORF72-linked ALS, we show that activation of the DDR precedes the appearance of alterations in mitochondrial functions. Importantly, mitochondrial parameters can be ameliorated by the inhibition of ATM and nicotinamide riboside supplementation. Overall, our data indicate the existence of a pathogenic crosstalk in which nuclear DNA damage promotes MD, which in turn may increase ROS production and DNA lesions. Considering that depletion of several ALS-linked proteins (such as TDP-43 and FUS) (Wang et al., 2021) can also lead to the accumulation of DNA damage, our study uncovers a potential common pathogenic mechanism between different ALS/FTD subtypes.

## Results

### DSBs in nuclear DNA induce deficits in mitochondrial function in AID-DIvA cells

Studies examining the DDR have generally been using ionizing radiation or treatments with genotoxic drugs to induce DNA damage in mammalian cells. These studies are inherently limited by the fact that these approaches affect both nuclear and mitochondrial DNA, as well as many multiple cellular components. To induce genomic DNA lesions while avoiding direct damage to mitochondria, we used the human cell line AID-DIvA (AID-AsiSI-ER-U2OS, Aymard et al., 2014). In these cells, the rare-cutting AsiSI restriction nuclease is expressed as a fusion protein with an auxin-inducible degron (AID) and the estrogen receptor (ER). Treatment with tamoxifen (4-OHT) triggers nuclear localization of the AsiSI enzyme and the rapid induction of sequence-specific DSBs at ∼100 annotated positions across the human genome (Iacovoni et al., 2010; Massip et al., 2010). The nuclease is also fused to an AID, which promotes rapid degradation of the enzyme upon auxin addition, allowing the study of DSB repair kinetics. Thus, in contrast to ionizing radiation or radiomimetic drugs, AID-DIvA cells allow the controlled introduction of DSBs exclusively in nuclear DNA (Supp. Figure 1A).

**Figure 1.**
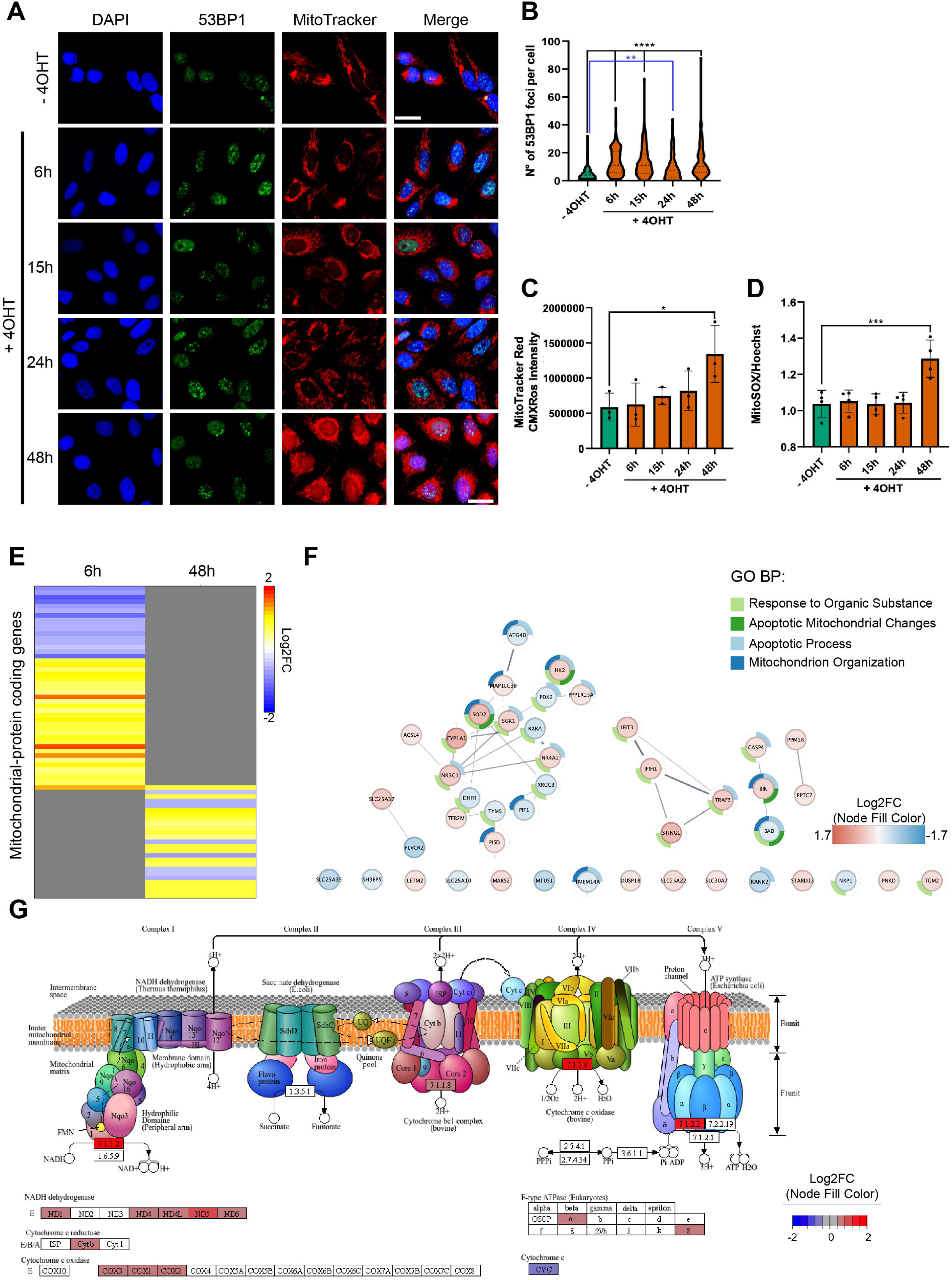
Nuclear DNA double-strand breaks induce mitochondrial dysfunction in AID-DIvA cells. A. Representative immunofluorescence images of AID-DIvA cells treated with 4-hydroxytamoxifen (4-OHT) for 6, 15, 24, and 48 h to induce AsiSI-mediated nuclear DNA double-strand breaks (DSBs). Cells were stained with anti-53BP1 antibodies and with MitoTracker Red to assess mitochondrial membrane potential (MMP). Nuclei were counterstained with DAPI. Scale bar, 20 μm. B. Quantification of 53BP1 foci in AID-DIvA cells following 4-OHT treatment for the indicated times. C. Quantification of MitoTracker Red fluorescence intensity in AID-DIvA cells treated with 4-OHT for the indicated times. D. Quantification of mitochondrial reactive oxygen species (ROS) production measured by MitoSOX Red fluorescence in AID-DIvA cells following DSB induction. E. Gene Ontology biological process enrichment analysis of mitochondrial differentially expressed genes (mtDEGs) identified by RNA-seq after 6 h and 48 h of DSB induction. F. Network representation of enriched biological processes associated with mtDEGs after 6 h of DSB induction. Node color indicates Log2 fold change (Log2FC), while node’s border color indicates the enriched process. G. Graphic representation of enriched OXPHOS biological processes associated with mtDEGs after 48 h of DSB induction. Node color indicates Log2 fold change (Log2FC). Data are presented as means ± SEM. Statistical significance was determined using one-way ANOVA followed by appropriate post hoc tests. *P < 0.05, **P < 0.01, ***P < 0.001, ****P < 0.0001.

To characterize the nuclear and mitochondrial damage timeline, we treated AID-DIvA cells with 4-OHT for 6, 15, 24, and 48 hours. Cells were stained with anti-53BP1 antibodies to monitor DNA repair foci, and with MitoTracker Red to assess mitochondrial membrane potential (MMP). As expected, 4-OHT treatment induced an increase in 53BP1 foci within 6 hours, which remained stable throughout the time-course (Figure 1A, B). In contrast, staining with MitoTracker Red revealed a significant increase in MMP only after 48 hours of 4-OHT treatment, indicative of mitochondrial hyperpolarization (Figure 1A, C). Consistent with this, mitochondrial reactive oxygen species (ROS) production measured by MitoSOX Red also showed a significant increase at the 48-hour time point (Figure 1 D). This pattern aligns with findings in other models where increased MMP accompanies enhanced mitochondrial ROS production, reflecting impaired electron transport chain efficiency and promoting oxidative stress (Heher et al., 2022; Xu et al., 2025). Together, these data support a model in which mitochondrial hyperpolarization and ROS accumulation occur sequentially during chronic nuclear DNA damage, contributing to progressive MD.

AsiSI-generated DSBs were shown to affect transcription both in the vicinity of the breaks as well as globally eliciting a DDR response that involves p53 (Iannelli et al., 2017). To investigate any specific change in the expression of mitochondrial genes in our experimental system, we profiled the transcriptome of uninduced, and induced DIvA cells at 6 and 48 hours by RNAseq. PCA analysis showed that the first two principal components (PCs) explained 54% of the variability among the samples. Both time points were grouped in distinct clusters well separated from the untreated control samples, indicating a clear difference in the transcriptomes (Supp. Figure 1B). Quantitative transcriptome differences between treated and untreated AID-DIvA cells were represented with two Volcano plots, revealing a total of 1,091 differentially expressed genes (DEGs) at 6 hours and 462 DEGs at 48 hours of DSBs induction, with selected transcripts encoding for mitochondrial proteins highlighted in both plots (Supp. Figure 1C, D). The intersection of the identified DEGs with the UniProt list of mitochondrial genes identified one gene overlapping between the two-time point (ABAT), and 44 and 24 differentially expressed mitochondrial genes (mtDEGs) at 6 h and 48 h, respectively (Supp. Figure 1E and Supp. Table 1). Notably, ABAT, a key enzyme linking GABA metabolism to mitochondrial nucleoside salvage and mtDNA maintenance, was uniquely regulated over time, suggesting its involvement in the cellular response to sustained DNA damage and MD (Besse et al., 2015; Zhang et al., 2022).

Although most mtDEGs were upregulated at both time points, gene set enrichment analysis showed distinct functional signatures. While after 6 hours of DSB induction the biological process analysis revealed that the most significantly affected pathways are related to apoptosis (Figure 1E, F), at 48 hours mtDEGs are enriched in gene sets related to “oxidative phosphorylation” and affecting electron transport chain components (Figure 1E, G). These findings confirm that the chronic induction of DSBs in nuclear DNA perturbs mitochondrial gene expression and function.

We then decided to better characterize the type of mitochondrial damage induced by DDR induction by performing Seahorse extracellular flux analyses. We observed a significant impairment of mitochondrial function as early as 6 hours after 4-OHT treatment, with reductions in basal respiration, ATP-linked respiration and maximal respiratory capacity that persisted at later time points (Figure 2A, B). Importantly, these effects do not appear to be caused by 4-OHT itself, as treatment of parental U2OS cells, which do not express the inducible AsiSI nuclease, did not result in detectable changes in mitochondrial respiration (Supp. Figure 1F, G), indicating that the observed defects are specifically associated with DSBs induction.

**Figure 2.**
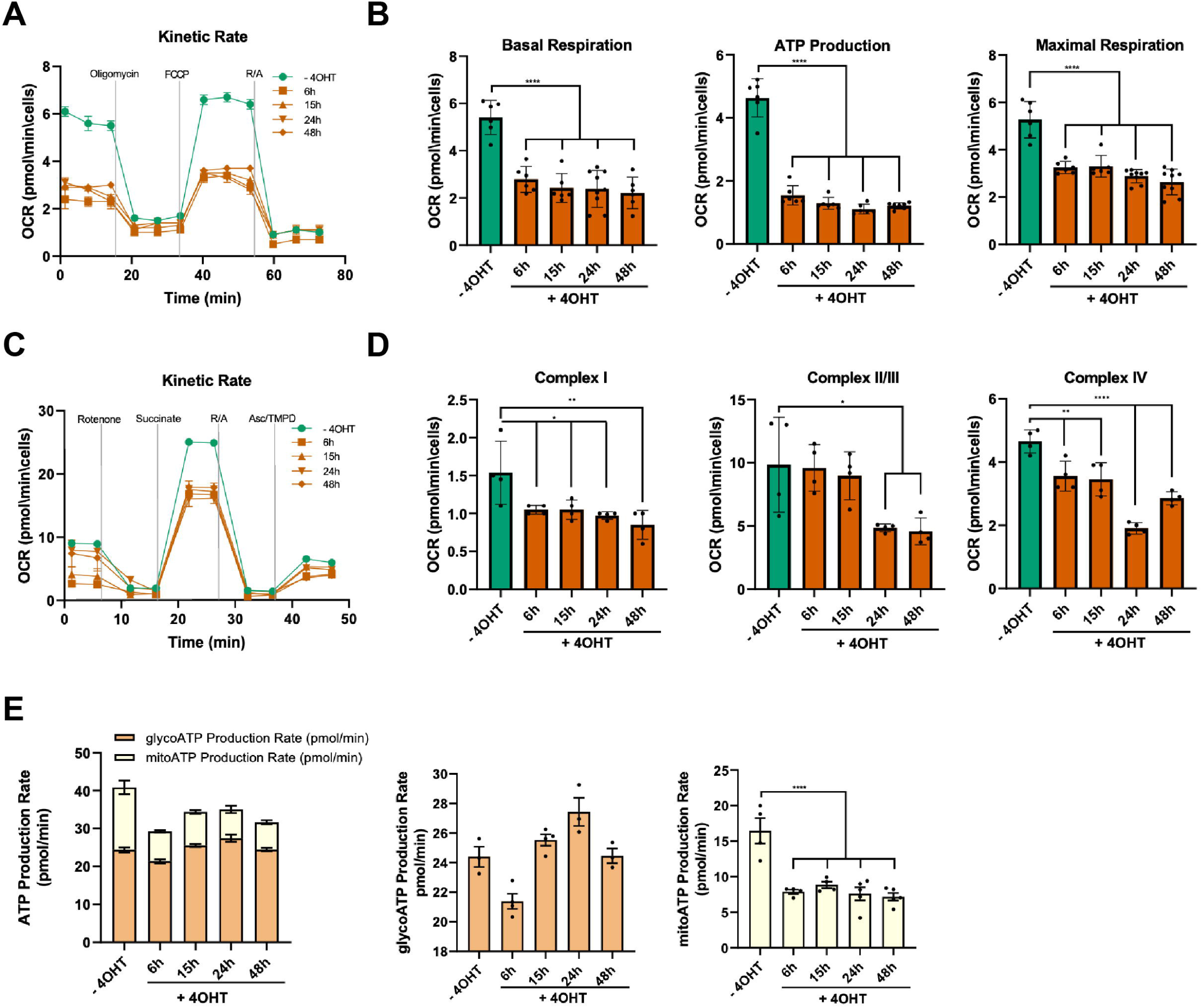
Nuclear DNA double-strand breaks impair mitochondrial respiration and electron transport chain activity. A. Representative Seahorse XF Cell Mito Stress Test profile of oxygen consumption rate (OCR) in AID-DIvA cells treated with 4-hydroxytamoxifen (4-OHT) for the indicated times to induce AsiSI-mediated double-strand breaks (DSBs). B. Quantification of basal respiration, ATP-linked respiration, and maximal respiratory capacity obtained from Seahorse XF Cell Mito Stress Test analyses in AID-DIvA cells following DSB induction. C. Representative Seahorse XF Electron Flow Assay profile used to evaluate the activity of individual electron transport chain (ETC) complexes in AID-DIvA cells treated with 4-OHT for the indicated times. D. Quantification of Complex I, Complex II/III, and Complex IV activities derived from Seahorse XF Electron Flow Assay analyses in AID-DIvA cells following DSB induction. E. Quantification of mitochondrial and glycolytic ATP production rates obtained by Seahorse XF Real-Time ATP Rate Assay in AID-DIvA cells following DSB induction. Data are presented as means ± SEM. Statistical significance was determined using one-way ANOVA followed by appropriate post hoc tests. *P < 0.05, **P < 0.01, ****P < 0.0001.

To dissect the origin of this impairment, we analyzed the activity of individual ETC complexes where we detected an early decrease in Complex I and Complex IV activity at 6 hours (Figure 2C, D), while Complex II/III activity was affected only at later time points, suggesting a progressive spread of dysfunction across the ETC rather than simultaneous impairment of all complexes. The early sensitivity of Complexes I and IV may reflect their dual genetic control by nuclear and mitochondrial genomes, making them particularly vulnerable to DDR-mediated nucleus-to-mitochondria signalling. A detailed analysis of ATP production revealed that DSB induction selectively compromises mitochondrial ATP generation while leaving glycolytic ATP production largely unaffected (Figure 2E), indicating that oxidative phosphorylation is specifically impaired without a compensatory increase in glycolysis.

Overall, our data suggest that nuclear DNA damage induces an early mitochondrial dysfunction at the bioenergetic level, whereas more pronounced alterations, such as membrane hyperpolarization and ROS production, emerge only following persistent stress, reflecting a progression from an initial adaptive phase to overt mitochondrial damage.

### Resolution of nuclear DNA DSBs restores mitochondrial function in AID-DIvA cells

To further confirm that MD is consequential to DNA DSB formation we performed a recovery experiment. After 48h of DSBs induction, AID-DIvA cells were treated with auxin to stop AsiSI activity. Auxin treatment induced degradation of the AID-AsiSi-ER fusion protein after 4 and 24 hours (Supplementary Figure 2A, B), and concomitantly a decrease of 53BP1 foci reflecting the progressive reduction of DSBs (Figure 3 A, B). Interestingly, we also observed a full recovery of all previously altered mitochondrial functions after 24 hours of auxin treatment (Figure 3C, D). Consistently, analysis of individual ETC complex activities revealed a robust recovery of Complexes I and IV activity, together with a clear trend toward restoration of Complex II/III function following 4-OHT treatment (Supp. Figure 2C, D). In line with these observations, a more detailed assessment of cellular ATP production confirmed that only mitochondrial ATP generation had been impaired by DSBs induction, whereas glycolytic ATP production remained unaffected throughout the experiment (Supp. Figure 2E), further supporting the specificity of the nuclear DNA damage-induced mitochondrial dysfunction.

**Figure 3.**
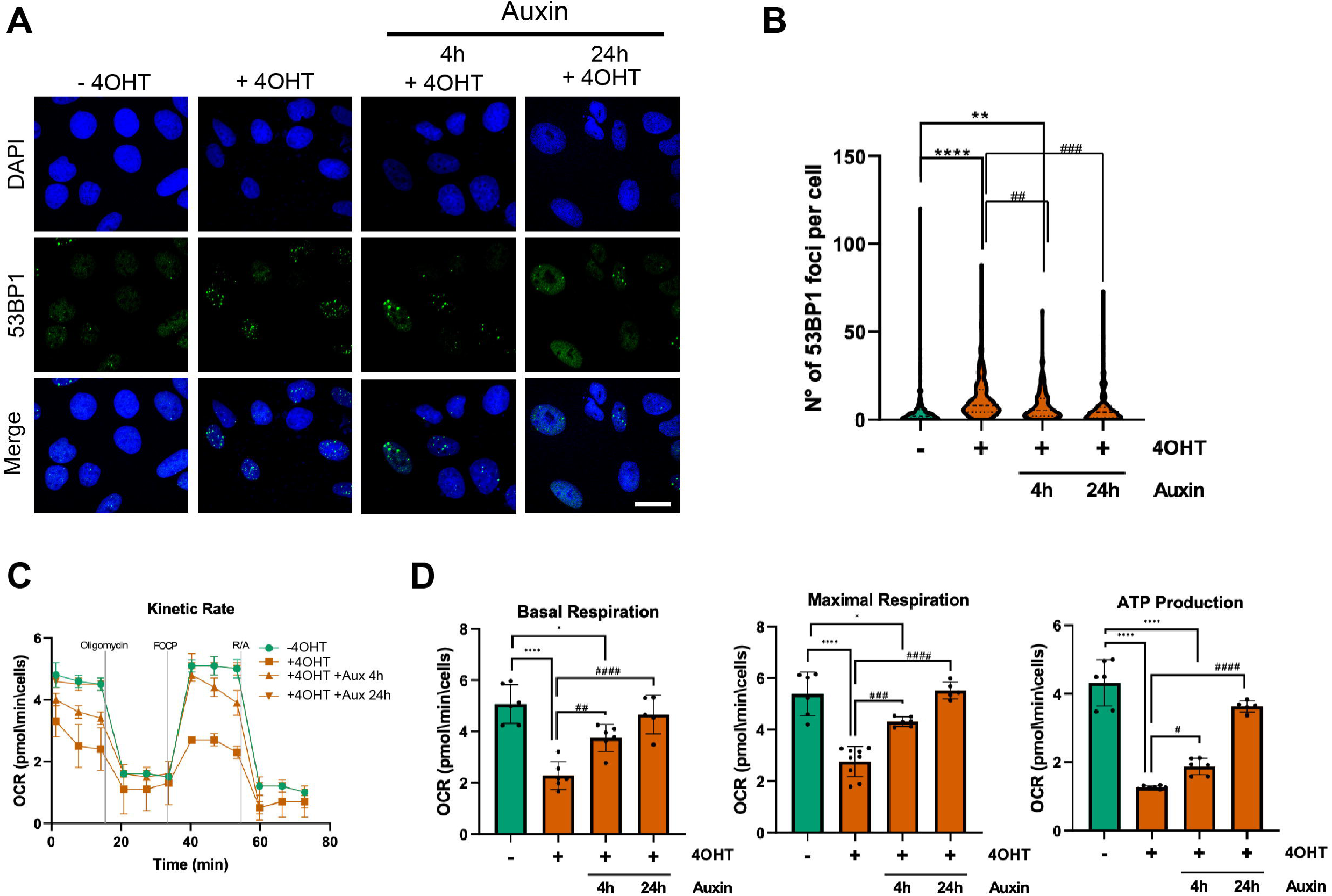
Resolution of nuclear DNA double-strand breaks restores mitochondrial function in AID-DIvA cells. A. Representative immunofluorescence images of 53BP1 foci in AID-DIvA cells following induction of AsiSI-mediated DNA double-strand breaks (DSBs) with 4-hydroxytamoxifen (4-OHT) and subsequent recovery after auxin treatment for the indicated times. Nuclei were counterstained with DAPI. Scale bar, 20 μm. B. Quantification of 53BP1 foci per nucleus during DSB induction and recovery upon auxin-mediated degradation of AsiSI. C. Representative Seahorse XF Cell Mito Stress Test profile of oxygen consumption rate (OCR) measured following DSB induction and recovery. D. Quantification of basal respiration, ATP-linked respiration, and maximal respiratory capacity derived from Seahorse XF Cell Mito Stress Test analyses during DSB recovery. Statistical significance was determined using one-way ANOVA followed by appropriate post hoc tests. *P < 0.05, **P < 0.01, ****P < 0.0001 compared with untreated; ^#^P < 0.05, ^##^P < 0.01, ^###^P < 0.001, ^####^P < 0.000.1 compared with 4OHT treated.

PARP1 activation represents one of the earliest steps in the DDR. PARP1 catalyzes the formation of poly(ADP-ribose) (PAR) polymers on target proteins using NAD+ as substrate, thereby initiating DNA repair signaling (Pandey & Black, 2021). In line with DSB induction, we observed a marked increase in poly(ADP-ribosyl)ated proteins 3 and 6 hours of 4-OHT treatment, and a strong reduction after 48 hours (Supp. Figure 3A). This process however consumes large amounts of NAD+, causing its rapid depletion, which has been shown to impair overall cellular metabolism (Alano et al., 2010). Consistent with increased parylation, assessment of NAD+ levels revealed a significant and persistent decrease in DIvA cells throughout the time course following 4-OHT treatment (Supp. Figure 3B). These findings suggest that PARP1-dependent NAD+ depletion contributes to the mitochondrial dysfunction observed upon DSB induction.

Based on these observations, we tested whether PARP1 inhibition with PJ34 could restore mitochondrial parameters in induced AID-DIvA cells. As shown in Figure 4A and 4B, treatment of induced DIvA cells for 4 h or 24h with the PARP1 inhibitor restored mitochondrial function increasing basal respiration, maximal respiratory capacity and ATP-linked respiration to control levels. Similarly, analysis of ATP production revealed that inhibition of PARP1 function selectively rescued mitochondrial ATP generation while leaving glycolytic ATP production largely unaffected (Supp. Figure 3C).

**Figure 4.**
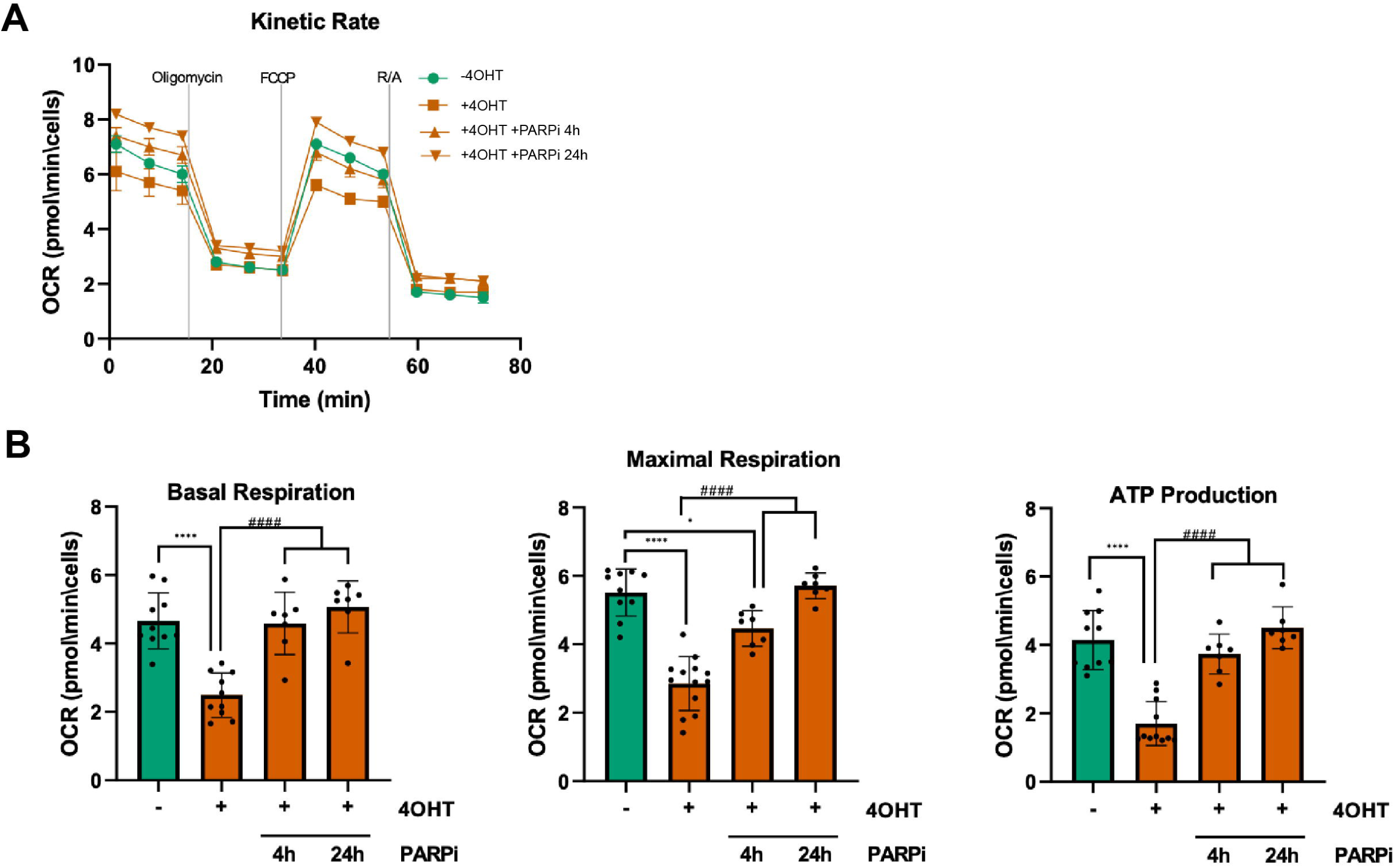
PARP inhibition rescues mitochondrial dysfunction induced by nuclear DNA double-strand breaks. A. Representative Seahorse XF Cell Mito Stress Test profile of oxygen consumption rate (OCR) in AID-DIvA cells treated with 4-OHT in the presence or absence of PJ34. B. Quantification of basal respiration, maximal respiratory capacity and ATP-linked respiration derived from Seahorse XF Cell Mito Stress Test analyses following PARP inhibition. Data are presented as means ± SEM. Statistical significance was determined using one-way ANOVA followed by appropriate post hoc tests. *P < 0.05, ****P < 0.0001 compared with untreated; ^####^P < 0.000.1 compared with 4OHT treated.

Collectively, these findings support a model in which DNA DSBs impair mitochondrial function via PARP1-mediated NADc depletion, with mitochondrial activity restored upon PARP inhibition or DSB resolution.

### Expression of (G4C2) repeats cause DDR activation and mitochondrial dysfunction

To determine if our observations were relevant for ALS pathology, we exploited a cell model of C9ORF72-ALS. Expansion of a GGGGCC (G4C2) repeat from 2-22 copies to 700-1600 copies in the first intron of the C9ORF72 gene is associated with 9p-linked ALS and in the related disorder frontotemporal dementia (FTD) (DeJesus-Hernandez et al., 2011; Renton et al., 2011). Studies showed that the expansions-containing RNA reduces C9ORF72 protein levels, can sequester RNA-binding proteins forming nuclear foci (Česnik et al., 2019), and can give rise to dipeptide repeats (DPRs) proteins via an unconventional repeat-associated non-AUG (RAN) translation mechanism leading to the accumulation of insoluble DPR aggregates (Ash et al., 2013; Mori et al., 2013). We chose this mutation because (i) the (G4C2) hexanucleotide expansion in *C9ORF72* represents the most common genetic alteration in fALS and in FTD; (ii) C9ORF72 patient-derived cells have altered mitochondrial bioenergetic function (Mehta et al., 2021; Onesto et al., 2016); (iii) it has been reported that markers of DNA damage are upregulated in motor neurons of C9ORF72-ALS patients in comparison to control subjects (Mehta et al., 2021), and that C9ORF72 DPRs inhibit DSB repair pathways that are utilized by post-mitotic neurons to maintain genomic integrity (Andrade et al., 2020; Lee et al., 2016).

We used a stable motor neuron-like NSC-34 cell model with tetracycline-inducible expression of 0 (sham) or 102 (G4C2) repeats (Stopford et al, 2017). NSC-34(G4C2)102 cells recapitulate both repeat RNA and DPR protein gain-of-function mechanisms previously described in multiple cellular and animal models of C9ORF72-ALS (Balendra & Isaacs, n.d.), both of which have been independently linked to DNA damage and mitochondrial dysfunction (Lopez-Gonzalez et al., 2016).

We first characterized NSC-34 sham and (G4C2)102 cells following tetracycline (Tet) induction for 2, 4, 6, and 7 days to verify that they recapitulate pathological features of cells derived from C9ORF72 ALS patients. We confirmed the increasing accumulation of characteristic sense (G4C2) RNA foci in the nucleus of (G4C2)102 cells, starting from day 2 of induction (Supp. Figure 4A, B), and of poly-GP DPR in Tet-induced NSC-34 (G4C2)102 cells. Poly-GP DPR levels were clearly detectable from induction-day 4 and remained uniformly expressed until the end of the time-course (Supp. Figure 4 C, D). Importantly, we did not detect any accumulation of poly-GR and poly-GP DPR in the mitochondrial fraction of (G4C2)102 induced-cells, suggesting that these DPRs accumulate just at cytoplasmic levels (Supp. Figure 4E). As expected, accumulation of nuclear RNA foci and cytoplasmic DPRs caused a decrease in cell viability in tetracycline-induced (G4C2)102 cells, particularly after 7 days of induction (Supp. Figure 4F).

Next, we investigated the timeline of DDR activation and mitochondrial damage in induced cells. Immunofluorescence experiments showed a significant increase in phosphorylated H2AX (γH2AX) and 53BP1 DNA damage foci in (G4C2)102-expressing cells, which started at induction day 2 and remained stable throughout the time-course (Figure 5 A-C). Conversely, analysis of MitoTracker Red CMXRos fluorescence in (G4C2)102 expressing cells showed a significant decrease in mitochondrial membrane potential only after 6 days of Tet-induction (Figure 5D, E), which was not matched by any alteration of mitochondrial ROS production (Figure 5F).

**Figure 5.**
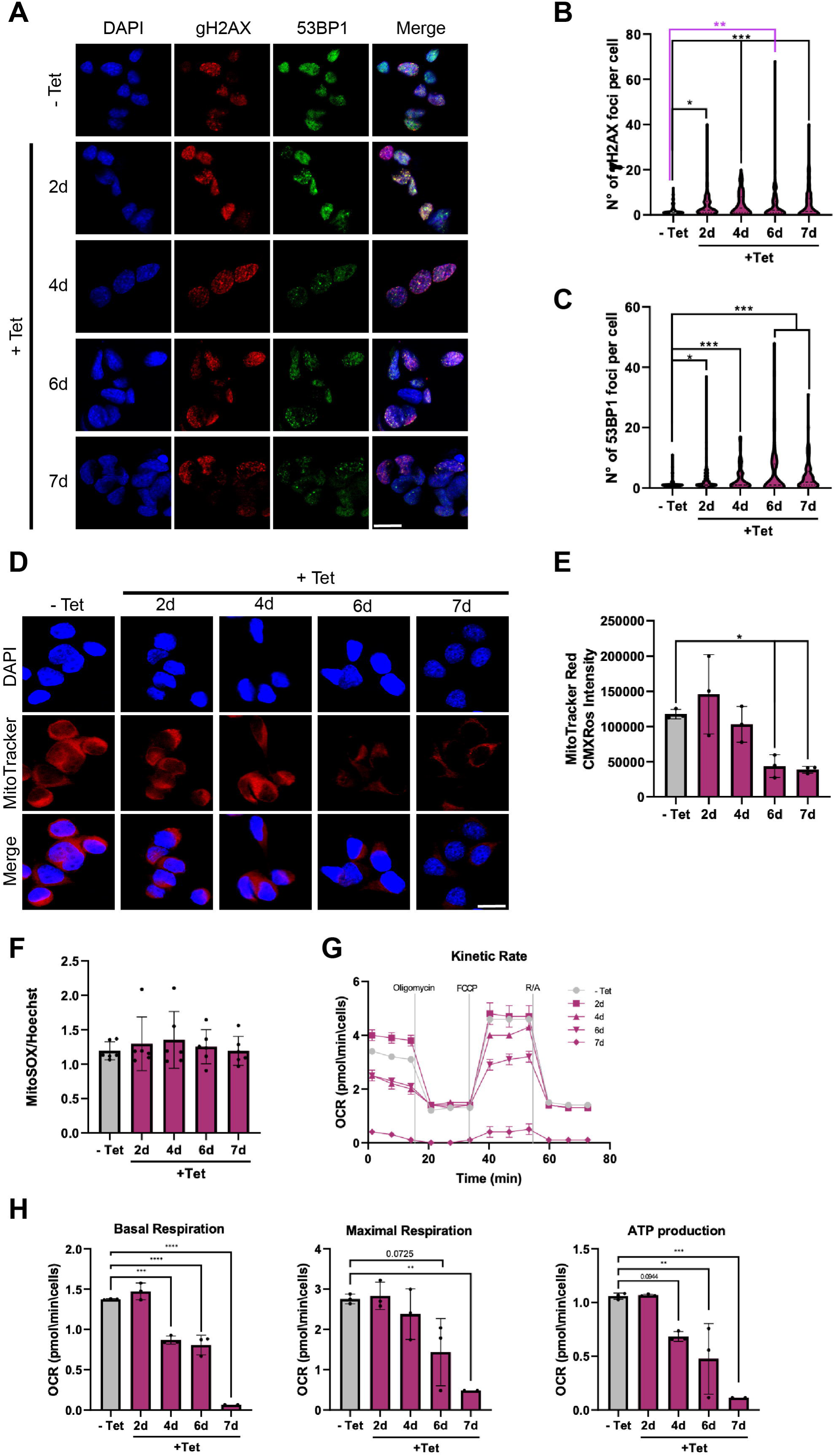
Expression of expanded (G4C2)102 repeats induces DDR activation and mitochondrial dysfunction in NSC-34 cells. A. Representative immunofluorescence images of NSC-34(G4C2)102 cells following tetracycline (Tet) induction for the indicated times. Cells were stained with anti-γH2AX and anti-53BP1 antibodies to assess DNA damage foci formation. Nuclei were counterstained with DAPI. Scale bar, 20 μm. B. Quantification of γH2AX-positive foci in NSC-34(G4C2)102 cells following Tet induction. C. Quantification of 53BP1-positive foci in NSC-34(G4C2)102 cells following Tet induction. D. Representative fluorescence images of NSC-34(G4C2)102 cells stained with MitoTracker Red CMXRos following Tet induction for the indicated times. Scale bar, 20 μm. E. Quantification of MitoTracker Red CMXRos fluorescence intensity in NSC-34(G4C2)102 cells following Tet induction. F. Quantification of mitochondrial reactive oxygen species (ROS) production measured by MitoSOX Red fluorescence in NSC-34(G4C2)102 cells following Tet induction. G. Representative Seahorse XF Cell Mito Stress Test traces of oxygen consumption rate (OCR) in NSC-34(G4C2)102 cells following Tet induction for the indicated times. H. Quantification of basal respiration and maximal respiratory capacity obtained from Seahorse XF Cell Mito Stress Test analyses in NSC-34(G4C2)102 cells following Tet induction. Data are presented as means ± SEM. Statistical significance was determined using one-way ANOVA followed by appropriate post hoc tests. *P < 0.05, **P < 0.01, ***P < 0.001, ****P < 0.0001.

These results suggest that mitochondrial dysfunction occurs after DDR activation, and DNA damage is not caused by an increase in mitochondrial ROS production.

In line with these findings, Seahorse extracellular flux analysis demonstrated a progressive decline in mitochondrial respiratory function in (G4C2)102 cells. Basal respiration was markedly reduced at later stages of induction, whereas maximal respiratory capacity showed a consistent downward trend across the time-course and reached significance at the latest time point examined (Figure 5G, H). Similarly, mitochondrial ATP production progressively declined and became significantly impaired at advanced stages, with an early tendency already detectable at intermediate time points (Supp. Figure 5A). In contrast, glycolytic ATP production exhibited only a modest, non-significant decrease, indicating that the energetic deficit was predominantly associated with mitochondrial dysfunction rather than with a generalized impairment of cellular ATP synthesis (Supp. Figure 5A).

The analysis of the activity of the respiratory chain complexes revealed an early functional impairment involving all complexes tested (Supp. Figure 5B, C). Importantly, sham cells subjected to the same Tet-induction protocol did not show any significant alteration in mitochondrial parameters, confirming that the observed defects were specifically associated with expression of expanded (G4C2) repeats (Supp. Figure 6).

**Figure 6.**
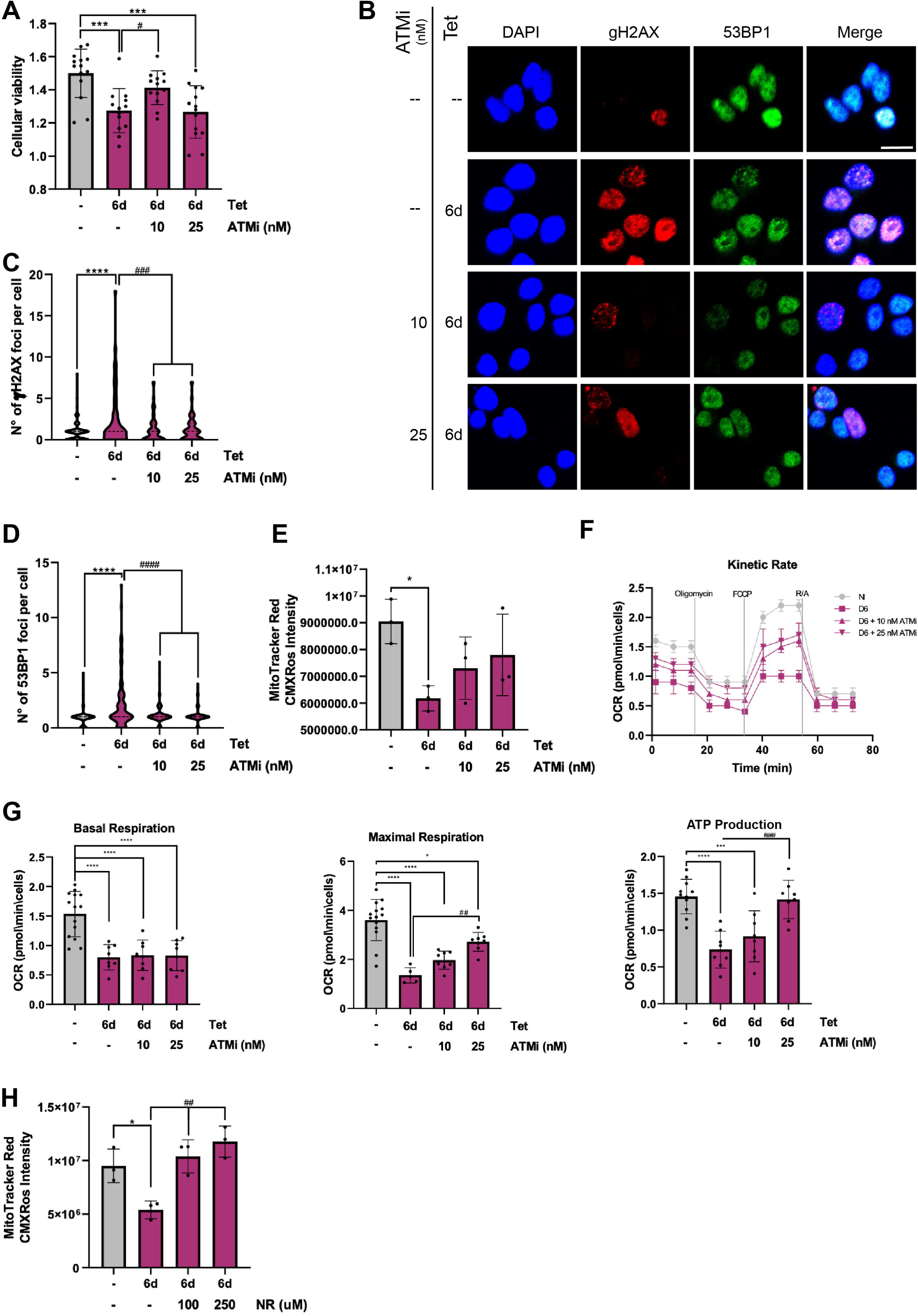
Inhibition of DDR signalling rescues mitochondrial dysfunction in NSC-34 (G4C2)102 cells. A. Cell viability analysis of NSC-34 (G4C2)102 cells following tetracycline (Tet)-induced expression of (G4C2) repeats and treatment with the ATM inhibitor KU-55933 (ATMi) at the indicated concentrations for 6 days. B. Representative immunofluorescence images of γH2AX and 53BP1 foci in NSC-34 (G4C2)102 cells following Tet induction in the presence or absence of ATMi treatment for 6 days. Nuclei were counterstained with DAPI. Scale bar, 20 μm. C. Quantification of γH2AX foci per nucleus in NSC-34 (G4C2)102. D. Quantification of 53BP1 foci per nucleus in NSC-34 (G4C2)102 cells. E. Quantification of mitochondrial membrane potential (MMP) in Tet-induced NSC-34 (G4C2)102 cells treated with KU-55933 for 6 days. F. Representative Seahorse XF Cell Mito Stress Test profile of oxygen consumption rate (OCR) in NSC-34 (G4C2)102 cells treated with the indicated concentrations of KU-55933. G. Quantification of basal respiration, maximal respiratory capacity and mitochondrial ATP production derived from Seahorse XF Cell Mito Stress Test analyses following ATM inhibition. H. Quantification of mitochondrial membrane potential in Tet-induced NSC-34 (G4C2)102 cells supplemented with Nicotinamide Riboside (NR) at the indicated concentrations for 6 days. Statistical significance was determined using one-way ANOVA followed by appropriate post hoc tests. *P < 0.05, ***P < 0.001, ****P < 0.0001 compared with untreated; ^#^P < 0.05, ^##^P < 0.01, ^###^P < 0.001 compared with 6 days Tet-treated.

Collectively, these results indicate that mitochondrial dysfunction develops progressively following expression of expanded (G4C2) repeats and becomes evident after the onset of DDR activation, supporting the view that mitochondrial impairment is a downstream consequence of early DNA damage signalling.

### DDR inhibition rescues mitochondrial function in the NSC34 (G4C2)102 cell model of C9ORF72-ALS

Since inhibition of DSBs induction in AID-DIvA cells rescued mitochondrial alterations, we decided to verify if suppression of DNA damage signalling could at least in part restore the mitochondrial damage in (G4C2)102-expressing NSC-34 cells. Of the three PIKK regulating the DDR, ATM responds primarily to DSBs, whereas ATR safeguards genomic integrity during S phase regulating the replication stress response. ATR is also activated by DSBs, however, through a mechanism that depends on the ATM (Jazayeri et al., 2006). Thus, to suppress DDR activation in (G4C2)102-expressing cells we used the ATM inhibitor KU-55933 (ATMi) at two sublethal concentrations that did not affect sham cells (data not shown). Interestingly, we observed that administration of 10 nM ATMi for six days resulted in a partial rescue of cell viability of (G4C2)102-expressing cells (Figure 6A). First, we tested the effect of the ATMi on DDR activation in cells expressing the (G4C2) repeats (Figure 6B-D). To quantify DDR activation, we measured the number of foci formed by γH2AX, which is a recognized indicator of DNA damage and a substrate of ATM, ATR, and DNA PKcs (Mah et al., 2010), and 53BP1. As expected, treatment with both ATMi concentrations for six days, concomitantly with the induction of (G4C2) repeats expression, drastically reduced the number of γH2AX and 53BP1 foci formed in (G4C2)102 cells when compared to untreated cells (Figure 6B-D). We also observed a partial rescue of the MMP in Tet-induced (G4C2)102 cells and co-treated with KU-55933 for 6 days (Figure 6E).

To further assess whether suppression of DDR signalling could restore mitochondrial bioenergetics in this model, we performed Seahorse extracellular flux analysis in Tet-induced NSC-34 sham control and in NSC-34 (G4C2)102 cells treated with the ATM inhibitor KU-55933. In agreement with the partial recovery of mitochondrial membrane potential (Figure 6E), pharmacological inhibition of ATM significantly improved mitochondrial respiratory performance. Although basal respiration remained largely unchanged, maximal respiration was markedly increased upon ATMi treatment in a dose-dependent manner (Figure 6F, G). Accordingly, while only a slight improvement was observed at the lower concentration, the effect became more pronounced at 25 nM KU-55933 reaching statistical significance (Figure 6F, G). Likewise, mitochondrial ATP production was significantly restored by ATMi, again showing a progressive response across concentrations and reaching near-normal levels at 25 nM (Figure 6F, G right panel). These findings were further supported by ATP rate assay measurements, which confirmed the recovery of mitochondrial ATP synthesis following ATM inhibition (Supp. Figure 7A). Collectively, these results indicate that attenuation of DDR signalling is sufficient to substantially rescue mitochondrial function in (G4C2)102-expressing cells, supporting a mechanistic link between persistent DNA damage signalling and mitochondrial dysfunction in this C9ORF72-ALS model.

Finally, based on the evidence that DDR induction in AID-DIvA cells led to a depletion of NAD+ (Supp. Figure 3B), we tested if boosting intracellular NAD(H) levels could have a beneficial effect on mitochondria. NSC-34 (G4C2)102 cells were concomitantly induced with tetracycline for 6 days and treated with 100 or 250 μM Nicotinamide Riboside (NR), a NAD+ precursor. As shown in Figure 6H, NR supplementation was sufficient to restore MMP. Overall, these data imply that boosting intracellular NAD(H) could have cytoprotective effects particularly in neurons that heavily depend on oxidative phosphorylation for energy production.

## Discussion

In this study, we provide evidence for a causal link between nuclear DNA damage and mitochondrial dysfunction, challenging the prevailing view that mitochondrial impairment is primarily the upstream driver of genomic instability in neurodegeneration. Using two complementary cellular models, we demonstrate that the induction of DSBs is sufficient to trigger mitochondrial dysfunction in the absence of direct mitochondrial genotoxic stress, and that this mechanism is relevant to C9ORF72-associated ALS.

In the AID-DIvA system, which allows the controlled generation of site-specific DSBs exclusively in nuclear DNA, we observed an early impairment of mitochondrial bioenergetics, characterized by reduced basal respiration, ATP-linked respiration and maximal respiratory capacity. Notably, these alterations occurred prior to overt mitochondrial damage, such as membrane hyperpolarization and increased ROS production, which became evident only upon prolonged DNA damage. This temporal sequence suggests that mitochondrial dysfunction is not an immediate consequence of oxidative stress, but rather an early downstream effect of nuclear DNA damage signaling. This observation is consistent with emerging evidence indicating that mitochondrial alterations can arise as a consequence of nuclear genome instability rather than being its primary cause (Farg et al., 2017).

Importantly, restoration of genome integrity through AsiSI degradation fully rescued mitochondrial function, demonstrating that the observed metabolic defects are reversible and directly dependent on the persistence of DNA lesions. This reversibility strengthens the causal relationship between nuclear DNA damage and mitochondrial dysfunction, and argues against non-specific effects of genotoxic treatments, which typically affect both nuclear and mitochondrial compartments.

Mechanistically, our data point to a central role of the DDR in mediating the crosstalk between the nucleus and mitochondria. Increasing evidence indicates that DDR signalling is tightly interconnected with cellular metabolism and mitochondrial function (Konopka & Atkin, 2022; Sun et al., 2020). In particular, we identified PARP1 activation as a key contributor to mitochondrial dysfunction. PARP1 hyperactivation upon DSB induction led to a significant depletion of cellular NAD+ levels, a critical cofactor for mitochondrial metabolism. This is in line with previous studies showing that excessive PARP1 activity consumes NAD+ and compromises cellular bioenergetics (Alano et al., 2010). Given the essential role of NAD+ in oxidative phosphorylation and redox balance, its depletion likely imposes a metabolic constraint on mitochondria, impairing ATP production and respiratory chain activity (Amjad et al., 2021).

Consistent with this model, pharmacological inhibition of PARP1 restored mitochondrial respiration and ATP synthesis, indicating that NAD+ consumption represents a major mechanistic link between DDR activation and mitochondrial dysfunction. While PARP1-dependent NAD+ depletion appears to play a prominent role, we cannot exclude the contribution of additional DDR-dependent signaling pathways, including ATM-mediated responses, which have also been implicated in metabolic regulation and mitochondrial homeostasis (Chow et al., 2019).

The relevance of this mechanism to ALS pathology is supported by our findings in a motor neuron-like model expressing expanded (G4C2) repeats of the C9ORF72 gene, the most common genetic cause of familial ALS (Balendra & Isaacs, n.d.). In these cells, DDR activation preceded the onset of mitochondrial dysfunction, mirroring the temporal dynamics observed in the AID-DIvA system. Importantly, mitochondrial defects developed progressively and were not associated with early increases in mitochondrial ROS, further supporting the notion that DNA damage is not secondary to oxidative stress in this context. Instead, our data indicate that persistent DDR signaling is sufficient to drive mitochondrial impairment.

These findings are consistent with previous reports showing that C9ORF72-associated pathological features, including DPR proteins and RNA foci, can induce DNA damage and impair DNA repair pathways (Andrade et al., 2020; Lee et al., 2016), as well as alter mitochondrial bioenergetics (Mehta et al., 2021; Onesto et al., 2016). Our results extend these observations by placing DDR activation upstream of mitochondrial dysfunction and identifying a mechanistic link between these two hallmarks of disease.

Consistent with this interpretation, pharmacological inhibition of ATM significantly ameliorated mitochondrial dysfunction in (G4C2)-expressing cells, restoring respiratory capacity and ATP production. These findings reinforce the idea that chronic activation of DDR pathways contributes directly to mitochondrial defects in ALS models. Moreover, boosting intracellular NAD(H) levels through supplementation with nicotinamide riboside improved both mitochondrial functions, further supporting the hypothesis that metabolic stress downstream of NAD+ depletion plays a key role in disease-associated toxicity (Yusri et al., 2025).

Our results integrate into a broader framework linking aging, genome instability, and mitochondrial dysfunction. Both increased DNA damage and impaired mitochondrial metabolism are well-established hallmarks of aging and neurodegenerative diseases, including ALS (Jagaraj et al., 2024; Smith et al., 2019). Traditionally, mitochondrial dysfunction has been proposed to promote DNA damage through increased ROS production and oxidative stress (Bogdanov et al., 2000). However, our findings support a bidirectional model in which nuclear DNA damage can act as an upstream driver of mitochondrial dysfunction via DDR-mediated metabolic reprogramming. This model is further supported by evidence from DNA repair-deficient disorders, where mitochondrial abnormalities arise as a consequence of genome instability (Farg et al., 2017).

Overall, our study identifies nuclear DNA damage as a primary trigger of mitochondrial dysfunction and highlights DDR signaling and NAD+ metabolism as key mediators of this process. These findings not only provide new insight into the pathogenic mechanisms underlying C9ORF72-associated ALS but also suggest that targeting DDR pathways or restoring NAD+ homeostasis may represent promising therapeutic strategies to counteract mitochondrial dysfunction and neurodegeneration.

## Materials & Methods

### Cell Lines, Cell Culture and Treatments

AID-DIvA cells were a kind gift of G. Legube (PMID: 32093728). AID-DIvA were cultured in high-glucose Dulbecco’s modified Eagle medium (DMEM) supplemented with 10% heat-inactivated fetal bovine serum (FBS), 100□U/mL penicillin, 100□µg/mL streptomycin, and 800 µg/ml G418. To induce the formation of AsiSI-dependent DNA double strand brakes (DSBs), AID-DIvA cells were treated with 300□nM 4-OHT for 6, 15, 24, or 48 hours. Conversely, addition of 500□µg/ml auxin for 4 or 24 hours allowed for AsiSI degradation in AID-DIvA cells previously treated with 300□nM 4-OHT for 48 hours, enabling DSB repair. To inhibit PARP, we treated AID-DIvA cells with 15nM PARP Inhibitor VIII, PJ34 for 4 or 24 hours following 48 hours 4-OHT treatment. For imaging experiments, AID-DIvA were plated onto 10mm coverslips. AID-DIvA were maintained at 37□°C and 5% CO2.

Sham control and (G4C2)102 NSC-34 cell lines were a kind gift of Pamela J. Shaw. NSC-34 cells were cultured in high-glucose Dulbecco’s modified Eagle medium (DMEM) supplemented with 10% tetracycline-free heat-inactivated FBS, 100□U/mL penicillin, 100□µg/mL streptomycin, 1 µg/mL blasticidin, and 50 µg/mL hygromycin B. Addition of 0.5 µg/mL tetracycline allowed for the expression of Sham control and (G4C2)102 plasmids for 2, 4, 6, or 7 days. Medium containing tetracycline was changed every two days to maintain the induction of the plasmids.

To inhibit ATM activity, tetracycline-induced (G4C2)102 cells were co-treated with 25 nM or 10 nM KU-55933 ATM inhibitor (SML1109) for 6 days. A similar approach was used for the Nicotinamide riboside (SMB00907, Merck) treatment. Indeed, tetracycline-induced (G4C2)102 cells were co-treated with 0.1 or 0.25 mM Nicotinamide riboside for 6 days and compared to non-induced cells.

For imaging experiments, NSC-34 cells were plated onto 10mm gelatin-coated coverslips. NSC-34 cells were maintained at 37□°C and 5% CO2.

### Transcriptomic and bioinformatic analysis

To perform the bulk transcriptomic experiments, total RNA was extracted from AID-DIvA cells using the RNeasy Plus Mini Kit (74134, QIAGEN) following manufacturer’s instructions. Briefly, cells were lysed in Buffer RLT Plus, and genomic DNA was eliminated from the lysates using the gDNA Eliminator spin column. Following three washes, RNA was eluted in RNase-free water supplemented with RNase inhibitors. Total RNA was quantified using a NanoDrop spectrophotometer (Thermo Fisher Scientific), and an equal amount of RNA was sent for sequencing using an Illumina HiSeq 2000 platform and performing a gene expression profiling using a ribosomal depletion approach. The analyzed samples included untreated controls (NT) and 4-OHT-treated cells collected after 6 h and 48 h of treatment, with two biological replicates per condition. The bioinformatic workflow started from demultiplexed paired-end FASTQ files. FASTQ files corresponding to the same biological sample were combined before quantification when required. Raw-read quality was assessed with FastQC (RRID:SCR_014583) and summarized with MultiQC (Ewels et al., 2016). Transcript abundance was estimated with kallisto (Bray et al., 2016) using 50 bootstrap iterations against a human transcriptome index generated from the Ensembl GRCh38 reference transcriptome (*Homo_sapiens.GRCh38.cdna.all.fa*), with gene annotation based on Ensembl release 112 (*Homo_sapiens.GRCh38.112.gtf*; Harrison et al., 2024). Quantification was performed in paired-end mode. Reads were quantified as unstranded. Transcript-level estimated counts were aggregated to gene-level counts using an Ensembl transcript-to-gene mapping.

Genes with low or background-level expression were removed using a sample-wise two-component Gaussian mixture filter applied to log2(1 + count) values; genes were retained if they were expressed in all replicates of at least one condition. Exploratory quality-control analyses included PCA/SVD-based sample clustering and inspection of sample-level expression structure. Differential expression analysis was then performed with DESeq2 (Love et al., 2014), using untreated cells as the reference condition and a design formula including treatment condition. Genes were considered differentially expressed when the Benjamini-Hochberg adjusted p-value was < 0.05 and the absolute log2 fold change was > 0.5.

Mitochondrial protein-coding genes were identified using UniProt subcellular-location annotations (Love et al., 2014) and intersected with the differentially expressed gene lists to define mitochondrial DEGs. Downstream analyses included PCA/sample clustering, volcano plots, log2 fold-change heatmaps, and gene-overlap plots. Gene Ontology over-representation analysis was performed with clusterProfiler (Yu et al., 2012) using org.Hs.eg.db annotations and Gene Ontology terms (Aleksander et al., 2023). Ensembl IDs were mapped to Entrez IDs, all GO ontologies were tested, p-values were adjusted using the Benjamini-Hochberg method, and enriched terms were retained using a q-value cutoff of 0.2 and a semantic-similarity simplification cutoff of 0.6. Terms with 10–500 overlapping input genes were retained. Mitochondrial DEG heatmap was built using Origin 2025b software, while network analysis was performed using the String app available at Cytoscape and the Pathviewer function available in R.

### Immunofluorescence

Following fixation with 4% PFA, permeabilization with 0.2% Triton x-100, and blocking with 5% FBS in PBS, cells were incubated with H2AX (sc-517348, Santa Cruz Biotechnology, 1:400 in 5% FBS diluted in PBS) and 53BP1 (4937S, Cell Signalling technology, 1:400 in 5% FBS diluted in PBS) primary antibodies overnight at 4°C. Cells were then incubated with Alexa Fluor Plus 488anti-rabbit and/or Alexa Fluor Plus 647 anti-mouse secondary antibodies for 1 hour at room temperature prior to staining them with DAPI for 5 minutes at room temperature. Coverslips were mounted on slides using a drop of mounting medium FluorSave Reagent (Millipore) and visualized through a Nikon A1R confocal optical confocal microscope. Nuclear H2AX and 53BP1 foci were calculated using the find maxima function within defined nuclear regions on ImageJ software, as previously described. The distribution of the number of spots in each nucleus, as well as the percentage of cells with a number of foci greater than one were visualized using violin plots. Unpaired t-test and one-way ANOVA were calculated using GraphPad Prism.

For MitoTracker Red CMXRos Staining AID-DIvA and NSC-34 cells were treated with 100nM MitoTracker Red CMXRos for 30 minutes at 37□°C and 5% CO2 before fixing them with 4% paraformaldehyde (PFA) for 20 minutes at room temperature and permeabilizing them with 0.2% Triton X-100 for 5 minutes at room temperature. NSC-34 cells were then directly stained with DAPI for 5 minutes at room temperature, while AID-DIvA were blocked with 5% FBS in PBS for 1 hour at room temperature before proceeding with the immunofluorescence protocol, as detailed below. The mitochondrial membrane potential of induced/treated vs non-induced/non-treated cells was calculated by measuring the raw integrated density of MitoTracker Red CMXRos intensity and dividing it by the number of nuclei in each field of view. Statistical analysis was performed by One Way ANOVA, using GraphPad Prism software.

### Bioenergetic analysis

Cellular bioenergetic profiling was assessed by extracellular flux analysis using a Seahorse XF96 Analyzer (Seahorse Bioscience–Agilent). XF96 microplates were coated with Cell-Tak (CLS354240 Corning), and NSC-34 motor neuron–like cells or DIvA cells were seeded at a density of 7,000 cells per well and allowed to adhere overnight. Bioenergetic measurements were performed in NSC-34 cells expressing either an empty vector (Sham) or a tetracycline-inducible expanded (G4C2)102 repeat following tetracycline induction for 2–7 days; in selected experiments, induction was restricted to 2 or 6 days and carried out in the presence or absence of the ATM inhibitor KU-55933 (10 or 25 nM) (SML1109 Sigma-Aldrich), with non-induced cells used as controls. In parallel, analyses were conducted in DIvA cells that were either left untreated or treated with tamoxifen alone or with tamoxifen in combination with auxin for 6, 15, 24, or 48 h, as indicated.

Immediately prior to extracellular flux measurements, culture medium was replaced with assay-specific Seahorse medium. For the Cell Mito Stress Test and the ATP Real-Time Rate Assay, culture medium was replaced with XF assay medium (Dulbecco’s Modified Eagle’s Medium, w/o glucose; Agilent Seahorse) supplemented with 1 mM sodium pyruvate, 10 mM glucose, and 2 mM L-glutamine, with the pH adjusted to 7.4. For assessment of mitochondrial electron transport chain activity, cells were incubated in respiration buffer containing 250 mM sucrose, 15 mM KCl, 1 mM EGTA, 5 mM MgCl□, and 30 mM K□HPO□, supplemented with 10 mM pyruvate, 2 mM malate, 4 mM ADP, and XF Plasma Membrane Permeabilizer (XF PMP, 1 nM). Prior to each assay, cells were equilibrated for 45 min at 37 °C in a CO□-free incubator.

Mitochondrial respiration was evaluated by monitoring the oxygen consumption rate (OCR) under basal conditions and following sequential injections of oligomycin (1.5 µM), carbonyl cyanide 4-(trifluoromethoxy) phenylhydrazone (FCCP, 1 µM), and a combination of rotenone (0.5 µM) and antimycin A (0.5 µM), allowing determination of basal respiration, ATP-linked respiration and maximal respiratory capacity. ATP production rates were determined using the ATP Real-Time Rate Assay by measuring OCR following injection of oligomycin (1.5 µM) and rotenone/antimycin A (0.5 µM each), enabling discrimination between mitochondrial and glycolytic ATP synthesis. Electron transport chain function was further assessed using the Electron Flow Assay, in which the XFe96 sensor cartridge was loaded with compounds to achieve final well concentrations of rotenone (2 µM), succinate (10 mM), antimycin A (4 µM), and a combination of ascorbate (10 mM) and N,N,N’,N’– tetramethyl-p-phenylenediamine (TMPD, 100 µM).

All Seahorse data were analyzed using Wave software (Agilent Technologies) and are presented as point-to-point OCR measurements normalized to cell number (pmol/min/cell). Cell number was determined for each well after completion of the assay by automated nuclei counting. Data are expressed as mean ± SEM, and statistical analyses were performed as indicated in the corresponding figure legends.

Intracellular NAD+ levels were quantified using the Invitrogen NAD+/NADH Colorimetric Assay Kit according to the manufacturer’s instructions. AID-DIvA cells were treated with 4-hydroxytamoxifen (4-OHT) for 6, 15, 24, and 48 h. Following treatment, cells were collected and lysed using the extraction buffer provided in the kit. Total NAD+ level was measured by a colorimetric assay based on WST-8 reduction, with absorbance read at 450 nm using a microplate reader.

### Cellular fractionation

Cytosolic and mitochondrial fractions were prepared from NSC-34 cells expressing either an empty vector (Sham) or a tetracycline-inducible expanded (G4C2)102 repeat. Cells were analyzed under non-induced (NI) conditions or following tetracycline induction for 2, 4, or 6 days. Cell pellets were homogenized in isolation buffer containing 210 mM mannitol, 70 mM sucrose, 1 mM EDTA, and 10 mM HEPES-KOH (pH 7.5) using a Potter–Elvehjem homogenizer equipped with a Teflon pestle, according to the subcellular fractionation protocol described by (Salvatori et al., 2018).

### Electrophoresis and Western blotting

Protein extracts obtained from total cell lysates or from cytosolic and mitochondrial fractions were separated by SDS–polyacrylamide gel electrophoresis and transferred onto nitrocellulose membranes (PerkinElmer). Membranes were blocked for 1 h in Tris-buffered saline containing 0.1% Tween-20 (TBS-T) supplemented with 5% non-fat dry milk, and subsequently incubated with primary antibodies Anti-C9ORF72/C9RANT(Poly-GP sense/antisense) (ABN1358 EMD Millipore, 1:2000), HA (3724 Cell Signaling, 1:1000), Poly/Mono-ADP Ribose (89190T Cell Signaling, 1:1000), B-Actin HRP-conjugated (sc-47778, Santa Cruz Biotechnology 1:7000), and B-Tubulin (sc-80005, Santa Cruz Biotechnology 1:7000) diluted in TBS-T containing 5% milk, either overnight at 4 °C or for 1 h at room temperature, as indicated.

Immunoreactive bands were visualized using enhanced chemiluminescence (Clarity™ Western ECL substrate; Bio-Rad). Apparent molecular weights were estimated by comparison with pre-stained molecular weight markers (Bio-Rad Laboratories).

Densitometric analysis was performed using ImageJ software (U.S. National Institutes of Health; RRID: SCR_003070). For subcellular fractionation experiments, the purity of cytosolic and mitochondrial fractions was assessed by immunoblotting for compartment-specific markers, and the relative abundance of proteins of interest was normalized to GAPDH (cytosolic marker) (Santa Cruz Biotechnology, Cat# sc-166545, RRID:AB_2107299 WB 1: 5000) or VDAC1 (mitochondrial marker) (Santa Cruz Biotechnology, Inc., Heidelberg Germany, Cat# sc-8829, RRID:AB_2214801, WB 1: 5000). For total cell lysates, protein expression levels were normalized to β-actin and tubulin antibodies. Statistical analysis for HA signal intensities in DIvA cells treated with 4OHT and Auxin was performed by One Way ANOVA, using GraphPad Prism software.

### RNA Fluorescent In Situ Hybridization

NSC-34 cells plated onto 10mm coverslips and treated with 0.5 µg/mL tetracycline for 2, 4, 6, and 7 days were fixed with 4% PFA for 15 minutes at room temperature. Following 2 PBS washes, fixed cells were permeabilized with 70% ethanol overnight at –20°C. Cells were then washed twice with 2x saline-sodium citrate buffer (SSC, 0.3 M sodium chloride and 0.03 M sodium citrate, pH 7) and once with 10% formamide diluted in 2x SSC buffer. Coverslips were then incubated overnight at 37°C with 5 M/5Cy3/mGmGmCmCmCmCmGmGmCmCmCmCmGmGmCmCmCmCmGmGmCmCmCm C FISH probe diluted in hybridization buffer (100 mg/ml dextran sulfate and 10% formamide in 2X SCC buffer). Following two washes in 10% formamide diluted in 2x SSC buffer for 30 minutes at 37°C in the dark, cells were stained with DAPI and mounted on slides using a drop of mounting medium FluorSave Reagent (Millipore). Coverslips were then visualized using Nikon A1R confocal optical confocal microscope to quantify nuclear RNA foci, as described above for the nuclear DNA damage foci. Statistical analysis was performed by One Way ANOVA, using GraphPad Prism software.

### MTT assay

AID-DIvA and NSC-34 cells were plated onto 96-well plates to measure cell vitality through the MTT (3-(4,5-dimethylthiazolyl-2)-2,5-diphenyltetrazolium bromide) assay. Following cell treatments, each well was treated with 250μg/ml MTT diluted in DMEM phenol red-free. Following the addition of the MTT solution and incubation for 3 hours, media was removed and DMSO added. When the solutions turned purple, the absorbance values could be read at the spectrophotometer Victor X3 (PerkinElmer) at 570nm wavelength. Unpaired t-test was calculated using GraphPad Prism.

### Statistical analysis

Statistical analysis was performed using GraphPad Prism software. Appropriate statistical tests were used, as specified throughout the “Material and Methods” section. R software was used to analyze bulk RNA seq data.

## Acknowledgements

We thank Prof. G. Legube and Prof. P.J. Shaw for providing AID-DIvA and NSC-34 Sham control and (G4C2)102 cells, respectively. The work was funded by the PRIN2022 grant n. 20222KSN2N of Italy’s MUR (Ministero dell’Università e della Ricerca).

## References

1. Alano, C. C., Garnier, P., Ying, W., Higashi, Y., Kauppinen, T. M., & Swanson, R. A. (2010). NAD+ depletion is necessary and sufficient for poly(ADP-ribose) polymerase-1-mediated neuronal death. The Journal of Neurosciencel: The Official Journal of the Society for Neuroscience, 30(8), 2967–2978. 10.1523/JNEUROSCI.5552-09.2010

2. Aleksander, S. A., Balhoff, J., Carbon, S., Cherry, J. M., Drabkin, H. J., Ebert, D., Feuermann, M., Gaudet, P., Harris, N. L., Hill, D. P., Lee, R., Mi, H., Moxon, S., Mungall, C. J., Muruganugan, A., Mushayahama, T., Sternberg, P. W., Thomas, P. D., Van Auken, K., … Westerfield, M. (2023). The Gene Ontology knowledgebase in 2023. Genetics, 224(1). 10.1093/GENETICS/IYAD031

3. Amjad, S., Nisar, S., Bhat, A. A., Shah, A. R., Frenneaux, M. P., Fakhro, K., Haris, M., Reddy, R., Patay, Z., Baur, J., & Bagga, P. (2021). Role of NAD+ in regulating cellular and metabolic signaling pathways. Molecular Metabolism, 49. 10.1016/j.molmet.2021.101195

4. Andrade, N. S., Ramic, M., Esanov, R., Liu, W., Rybin, M. J., Gaidosh, G., Abdallah, A., Del’olio, S., Huff, T. C., Chee, N. T., Anatha, S., Gendron, T. F., Wahlestedt, C., Zhang, Y., Benatar, M., Mueller, C., & Zeier, Z. (2020). Dipeptide repeat proteins inhibit homology-directed DNA double strand break repair in C9ORF72 ALS/FTD. Molecular Neurodegeneration, 15(1). 10.1186/S13024-020-00365-9

5. Ash, P. E. A., Bieniek, K. F., Gendron, T. F., Caulfield, T., Lin, W. L., DeJesus-Hernandez, M., Van Blitterswijk, M. M., Jansen-West, K., Paul, J. W., Rademakers, R., Boylan, K. B., Dickson, D. W., & Petrucelli, L. (2013). Unconventional Translation of C9ORF72 GGGGCC Expansion Generates Insoluble Polypeptides Specific to c9FTD/ALS. Neuron, 77(4), 639–646. 10.1016/j.neuron.2013.02.004

6. Aymard, F., Bugler, B., Schmidt, C. K., Guillou, E., Caron, P., Briois, S., Iacovoni, J. S., Daburon, V., Miller, K. M., Jackson, S. P., & Legube, G. (2014). Transcriptionally active chromatin recruits homologous recombination at DNA double-strand breaks. Nature Structural & Molecular Biology, 21(4), 366–374. 10.1038/NSMB.2796

7. Balendra, R., & Isaacs, A. M. (n.d.). C9orf72-mediated ALS and FTD: multiple pathways to disease. 10.1038/s41582-018-0047-2

8. Balendra, R., & Isaacs, A. M. (2018). C9orf72-mediated ALS and FTD: multiple pathways to disease. Nature Reviews. Neurology, 14(9), 544–558. 10.1038/S41582-018-0047-2

9. Besse, A., Wu, P., Bruni, F., Donti, T., Graham, B. H., Craigen, W. J., McFarland, R., Moretti, P., Lalani, S., Scott, K. L., Taylor, R. W., & Bonnen, P. E. (2015). The GABA transaminase, ABAT, is essential for mitochondrial nucleoside metabolism. Cell Metabolism, 21(3), 417–427. 10.1016/J.CMET.2015.02.008

10. Bogdanov, M., Brown, R. H., Matson, W., Smart, R., Hayden, D., O’Donnell, H., Flint Beal, M., & Cudkowicz, M. (2000). Increased oxidative damage to DNA in ALS patients. Free Radical Biology and Medicine, 29(7), 652–658. 10.1016/S0891-5849(00)00349-X

11. Bray, N. L., Pimentel, H., Melsted, P., & Pachter, L. (2016). Near-optimal probabilistic RNA-seq quantification. Nature Biotechnology, 34(5), 525–527. 10.1038/NBT.3519

12. Česnik, A. B., Darovic, S., Mihevc, S. P., Štalekar, M., Malnar, M., Motaln, H., Lee, Y. B., Mazej, J., Pohleven, J., Grosch, M., Modic, M., Fonovič, M., Turk, B., Drukker, M., Shaw, C. E., & Rogelj, B. (2019). Nuclear RNA foci from C9ORF72 expansion mutation form paraspeckle-like bodies. Journal of Cell Science, 132(5). 10.1242/JCS.224303

13. Chow, H. M., Cheng, A., Song, X., Swerdel, M. R., Hart, R. P., & Herrup, K. (2019). ATM is activated by ATP depletion and modulates mitochondrial function through NRF1. The Journal of Cell Biology, 218(3), 909–928. 10.1083/JCB.201806197

14. Cucchi, D., Gibson, A., & Martin, S. a. (2021). The emerging relationship between metabolism and DNA repair. *Cell Cycle (Georgetown*, Tex.), 20(10), 943–959. 10.1080/15384101.2021.1912889

15. DeJesus-Hernandez, M., Mackenzie, I. R., Boeve, B. F., Boxer, A. L., Baker, M., Rutherford, N. J., Nicholson, A. M., Finch, N. A., Gilmer, F., Adamson, J., Kouri, N., Wojtas, A., Sengdy, P., Hsiung, G.-Y. R., Karydas, A., Seeley, W. W., Josephs, K. A., Geschwind, D. H., Wszolek, Z. K., … Boylan, K. (2011). Expanded GGGGCC hexanucleotide repeat in non-coding region of C9ORF72 causes chromosome 9p-linked frontotemporal dementia and amyotrophic lateral sclerosis. Neuron, 72(2), 245–256. 10.1016/j.neuron.2011.09.011.Expanded

16. Ewels, P., Magnusson, M., Lundin, S., & Käller, M. (2016). MultiQC: summarize analysis results for multiple tools and samples in a single report. *Bioinformatics (Oxford*, England), 32(19), 3047–3048. 10.1093/BIOINFORMATICS/BTW354

17. Farg, M. A., Konopka, A., Soo, K. Y., Ito, D., & Atkin, J. D. (2017). The DNA damage response (DDR) is induced by the C9orf72 repeat expansion in amyotrophic lateral sclerosis. Human Molecular Genetics, 26(15), 2882–2896. 10.1093/hmg/ddx170

18. Harrison, P. W., Amode, M. R., Austine-Orimoloye, O., Azov, A. G., Barba, M., Barnes, I., Becker, A., Bennett, R., Berry, A., Bhai, J., Bhurji, S. K., Boddu, S., Lins, P. R. B., Brooks, L., Ramaraju, S. B., Campbell, L. I., Martinez, M. C., Charkhchi, M., Chougule, K., … Yates, A. D. (2024). Ensembl 2024. Nucleic Acids Research, 52(D1), D891–D899. 10.1093/NAR/GKAD1049

19. Heher, P., Ganassi, M., Weidinger, A., Engquist, E. N., Pruller, J., Nguyen, T. H., Tassin, A., Declèves, A. E., Mamchaoui, K., Banerji, C. R. S., Grillari, J., Kozlov, A. V., & Zammit, P. S. (2022). Interplay between mitochondrial reactive oxygen species, oxidative stress and hypoxic adaptation in facioscapulohumeral muscular dystrophy: Metabolic stress as potential therapeutic target. Redox Biology, 51. 10.1016/j.redox.2022.102251

20. Iacovoni, J. S., Caron, P., Lassadi, I., Nicolas, E., Massip, L., Trouche, D., & Legube, G. (2010). High-resolution profiling of gammaH2AX around DNA double strand breaks in the mammalian genome. The EMBO Journal, 29(8), 1446–1457. 10.1038/EMBOJ.2010.38

21. Iannelli, F., Galbiati, A., Capozzo, I., Nguyen, Q., Magnuson, B., Michelini, F., D’Alessandro, G., Cabrini, M., Roncador, M., Francia, S., Crosetto, N., Ljungman, M., Carninci, P., & D’Adda Di Fagagna, F. (2017). A damaged genome’s transcriptional landscape through multilayered expression profiling around in situ-mapped DNA double-strand breaks. Nature Communications, 8. 10.1038/NCOMMS15656

22. Jagaraj, C. J., Shadfar, S., Kashani, S. A., Saravanabavan, S., Farzana, F., & Atkin, J. D. (2024). Molecular hallmarks of ageing in amyotrophic lateral sclerosis. Cellular and Molecular Life Sciencesl: CMLS, 81(1). 10.1007/S00018-024-05164-9

23. Jazayeri, A., Falck, J., Lukas, C., Bartek, J., Smith, G. C. M., Lukas, J., & Jackson, S. P. (2006). ATM– and cell cycle-dependent regulation of ATR in response to DNA double-strand breaks. Nature Cell Biology, 8(1), 37–45. 10.1038/NCB1337

24. Konopka, A., & Atkin, J. D. (2022). DNA Damage, Defective DNA Repair, and Neurodegeneration in Amyotrophic Lateral Sclerosis. Frontiers in Aging Neuroscience, 14. 10.3389/FNAGI.2022.786420

25. Lee, K. H., Zhang, P., Kim, H. J., Mitrea, D. M., Sarkar, M., Freibaum, B. D., Cika, J., Coughlin, M., Messing, J., Molliex, A., Maxwell, B. A., Kim, N. C., Temirov, J., Moore, J., Kolaitis, R. M., Shaw, T. I., Bai, B., Peng, J., Kriwacki, R. W., & Taylor, J. P. (2016). C9orf72 Dipeptide Repeats Impair the Assembly, Dynamics, and Function of Membrane-Less Organelles. Cell, 167(3), 774–788.e17. 10.1016/j.cell.2016.10.002

26. Lopez-Gonzalez, R., Lu, Y., Gendron, T. F., Karydas, A., Tran, H., Yang, D., Petrucelli, L., Miller, B. L., Almeida, S., & Gao, F. B. (2016). Poly(GR) in C9ORF72-Related ALS/FTD Compromises Mitochondrial Function and Increases Oxidative Stress and DNA Damage in iPSC-Derived Motor Neurons. Neuron, 92(2), 383–391. 10.1016/J.NEURON.2016.09.015

27. Love, M. I., Huber, W., & Anders, S. (2014). Moderated estimation of fold change and dispersion for RNA-seq data with DESeq2. Genome Biology, 15(12). 10.1186/S13059-014-0550-8

28. Ma, W., & Zhou, S. (2025). Metabolic Rewiring in the Face of Genomic Assault: Integrating DNA Damage Response and Cellular Metabolism. Biomolecules, 15(2). 10.3390/BIOM15020168

29. Mah, L. J., El-Osta, A., & Karagiannis, T. C. (2010). gammaH2AX: a sensitive molecular marker of DNA damage and repair. Leukemia, 24(4), 679–686. 10.1038/LEU.2010.6

30. Massip, L., Caron, P., Iacovoni, J. S., Trouche, D., & Legube, G. (2010). Deciphering the chromatin landscape induced around DNA double strand breaks. *Cell Cycle (Georgetown*, Tex.), 9(15), 3035–3044. 10.4161/CC.9.15.12412

31. Mehta, A. R., Gregory, J. M., Dando, O., Carter, R. N., Burr, K., Nanda, J., Story, D., McDade, K., Smith, C., Morton, N. M., Mahad, D. J., Hardingham, G. E., Chandran, S., & Selvaraj, B. T. (2021). Mitochondrial bioenergetic deficits in C9orf72 amyotrophic lateral sclerosis motor neurons cause dysfunctional axonal homeostasis. Acta Neuropathologica, 141(2), 257–279. 10.1007/S00401-020-02252-5

32. Mori, K., Weng, S. M., Arzberger, T., May, S., Rentzsch, K., Kremmer, E., Schmid, B., Kretzschmar, H. A., Cruts, M., Van Broeckhoven, C., Haass, C., & Edbauer, D. (2013). The C9orf72 GGGGCC repeat is translated into aggregating dipeptide-repeat proteins in FTLD/ALS. *Science (New York*, N.Y.), 339(6125), 1335–1338. 10.1126/SCIENCE.1232927

33. Nijs, M., & Van Damme, P. (2024). The genetics of amyotrophic lateral sclerosis. Current Opinion in Neurology, 37(5), 560–569. 10.1097/WCO.0000000000001294

34. Onesto, E., Colombrita, C., Gumina, V., Borghi, M. O., Dusi, S., Doretti, A., Fagiolari, G., Invernizzi, F., Moggio, M., Tiranti, V., Silani, V., & Ratti, A. (2016). Gene-specific mitochondria dysfunctions in human TARDBP and C9ORF72 fibroblasts. Acta Neuropathologica Communications, 4(1), 47. 10.1186/S40478-016-0316-5

35. Pandey, N., & Black, B. E. (2021). Rapid Detection and Signaling of DNA Damage by PARP-1. Trends in Biochemical Sciences, 46(9), 744–757. 10.1016/j.tibs.2021.01.014

36. Renton, A. E., Majounie, E., Waite, A., Simón-Sánchez, J., Rollinson, S., Gibbs, J. R., Schymick, J. C., Laaksovirta, H., van Swieten, J. C., Myllykangas, L., Kalimo, H., Paetau, A., Abramzon, Y., Remes, A. M., Kaganovich, A., Scholz, S. W., Duckworth, J., Ding, J., Harmer, D. W., … Traynor, B. J. (2011). A hexanucleotide repeat expansion in C9ORF72 is the cause of chromosome 9p21-linked ALS-FTD. Neuron, 72(2), 257–268. 10.1016/J.NEURON.2011.09.010

37. Salvatori, I., Ferri, A., Scaricamazza, S., Giovannelli, I., Serrano, A., Rossi, S., D’Ambrosi, N., Cozzolino, M., Giulio, A. Di, Moreno, S., Valle, C., & Carrì, M. T. (2018). Differential toxicity of TAR DNA-binding protein 43 isoforms depends on their submitochondrial localization in neuronal cells. Journal of Neurochemistry, 146(5), 585–597. 10.1111/JNC.14465

38. Smith, E. F., Shaw, P. J., & De Vos, K. J. (2019). The role of mitochondria in amyotrophic lateral sclerosis. Neuroscience Letters, 710, 132933. 10.1016/j.neulet.2017.06.052

39. Sun, Y., Curle, A. J., Haider, A. M., & Balmus, G. (2020). The role of DNA damage response in amyotrophic lateral sclerosis. Essays in Biochemistry, 64(5), 847–861. 10.1042/EBC20200002

40. Tefera, T. W., & Borges, K. (2017). Metabolic Dysfunctions in Amyotrophic Lateral Sclerosis Pathogenesis and Potential Metabolic Treatments. Frontiers in Neuroscience, 10(JAN). 10.3389/FNINS.2016.00611

41. Wang, H., Kodavati, M., Britz, G. W., & Hegde, M. L. (2021). DNA Damage and Repair Deficiency in ALS/FTD-Associated Neurodegeneration: From Molecular Mechanisms to Therapeutic Implication. Frontiers in Molecular Neuroscience, 14. 10.3389/FNMOL.2021.784361

42. Xu, X., Pang, Y., & Fan, X. (2025). Mitochondria in oxidative stress, inflammation and aging: from mechanisms to therapeutic advances. Signal Transduction and Targeted Therapy, 10(1). 10.1038/S41392-025-02253-4

43. Yu, G., Wang, L. G., Han, Y., & He, Q. Y. (2012). clusterProfiler: an R package for comparing biological themes among gene clusters. Omicsl: A Journal of Integrative Biology, 16(5), 284–287. 10.1089/OMI.2011.0118

44. Yusri, K., Jose, S., Vermeulen, K. S., Tan, T. C. M., & Sorrentino, V. (2025). The role of NAD+ metabolism and its modulation of mitochondria in aging and disease. Npj Metabolic Health and Disease, 3(1). 10.1038/S44324-025-00067-0

45. Zhang, M., Zhong, H., Cao, T., Huang, Y., Ji, X., Fan, G. C., & Peng, T. (2022). Gamma-Aminobutyrate Transaminase Protects against Lipid Overload-Triggered Cardiac Injury in Mice. International Journal of Molecular Sciences, 23(4). 10.3390/IJMS23042182

