## Supplementary Figures for "Nuclear DNA Damage is a Primary Driver of Mitochondrial Dysfunction in *C9ORF72* ALS/FTD"

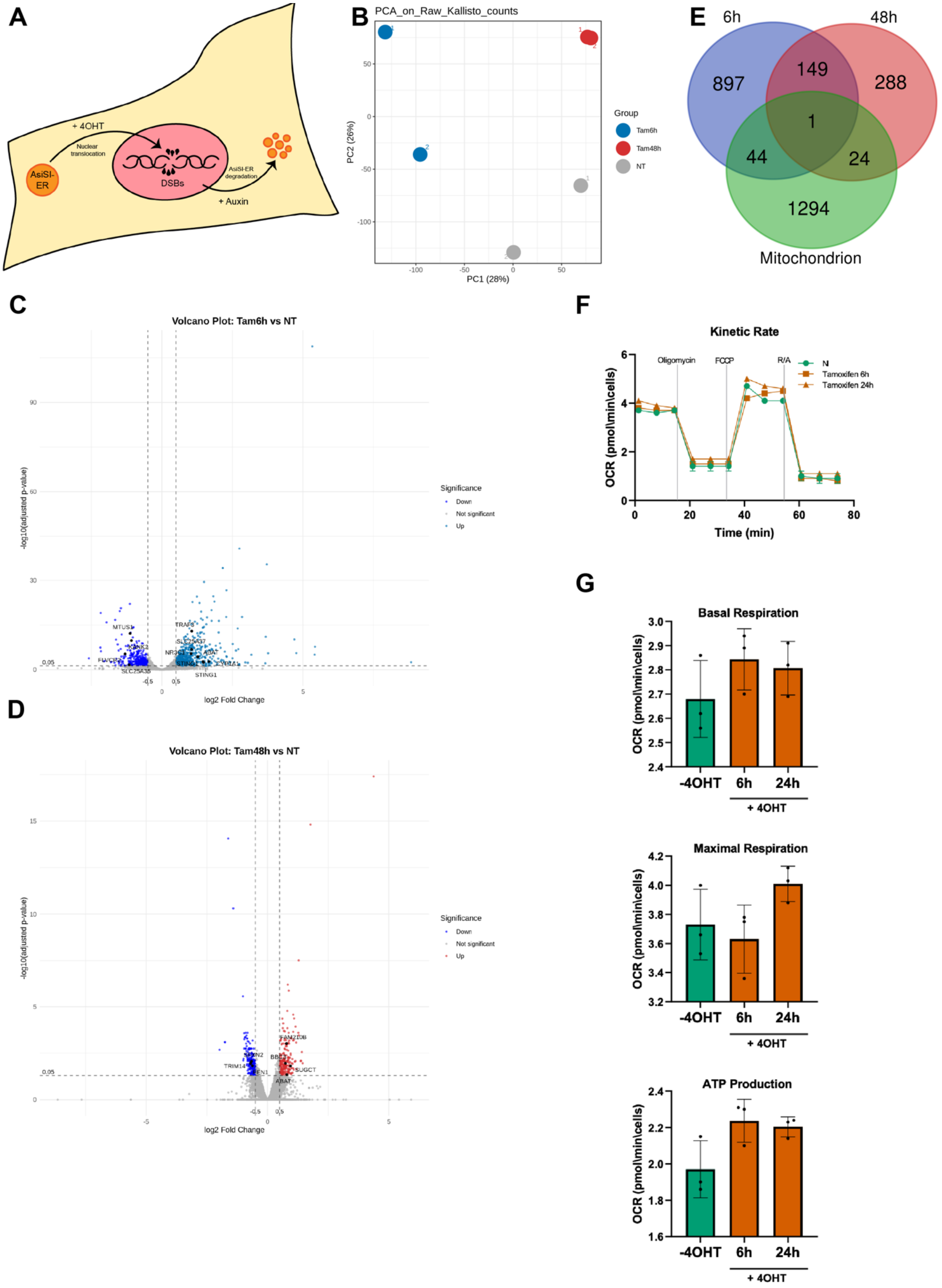


**Supplementary Figure 1**

**Supplementary Figure 1.**

A. Schematic representation of the AID-DIvA system used to induce sequence-specific double-strand breaks (DSBs) in nuclear DNA upon 4-hydroxytamoxifen (4-OHT) treatment.
B. Principal component analysis (PCA) of RNA-seq datasets from untreated AID-DIvA cells and cells untreated (NT) or treated with 4-OHT (Tam) for 6 h and 48 h.
C. Volcano plots showing differentially expressed genes (DEGs) identified by RNA-seq analysis after 6 h of DSB induction. Selected transcripts encoding mitochondrial proteins are highlighted.
D. Volcano plots showing differentially expressed genes (DEGs) identified by RNA-seq analysis after 48 h of DSB induction. Selected transcripts encoding mitochondrial proteins are highlighted.
E. Venn diagram showing the overlap between differentially expressed genes (mtDEGs, identified at 6 h and 48 h after DSB induction) and mitochondrial protein-coding genes (mitochondrion) identified in the UniProt database.
F. Representative Seahorse XF Cell Mito Stress Test profile of oxygen consumption rate (OCR) in parental U2OS cells treated with 4-OHT for the indicated times.
G. Quantification of basal respiration, ATP-linked respiration, and maximal respiratory capacity in parental U2OS cells treated with 4-OHT. Data are presented as means ± SEM. Statistical significance was determined using one-way ANOVA followed by appropriate post hoc tests.

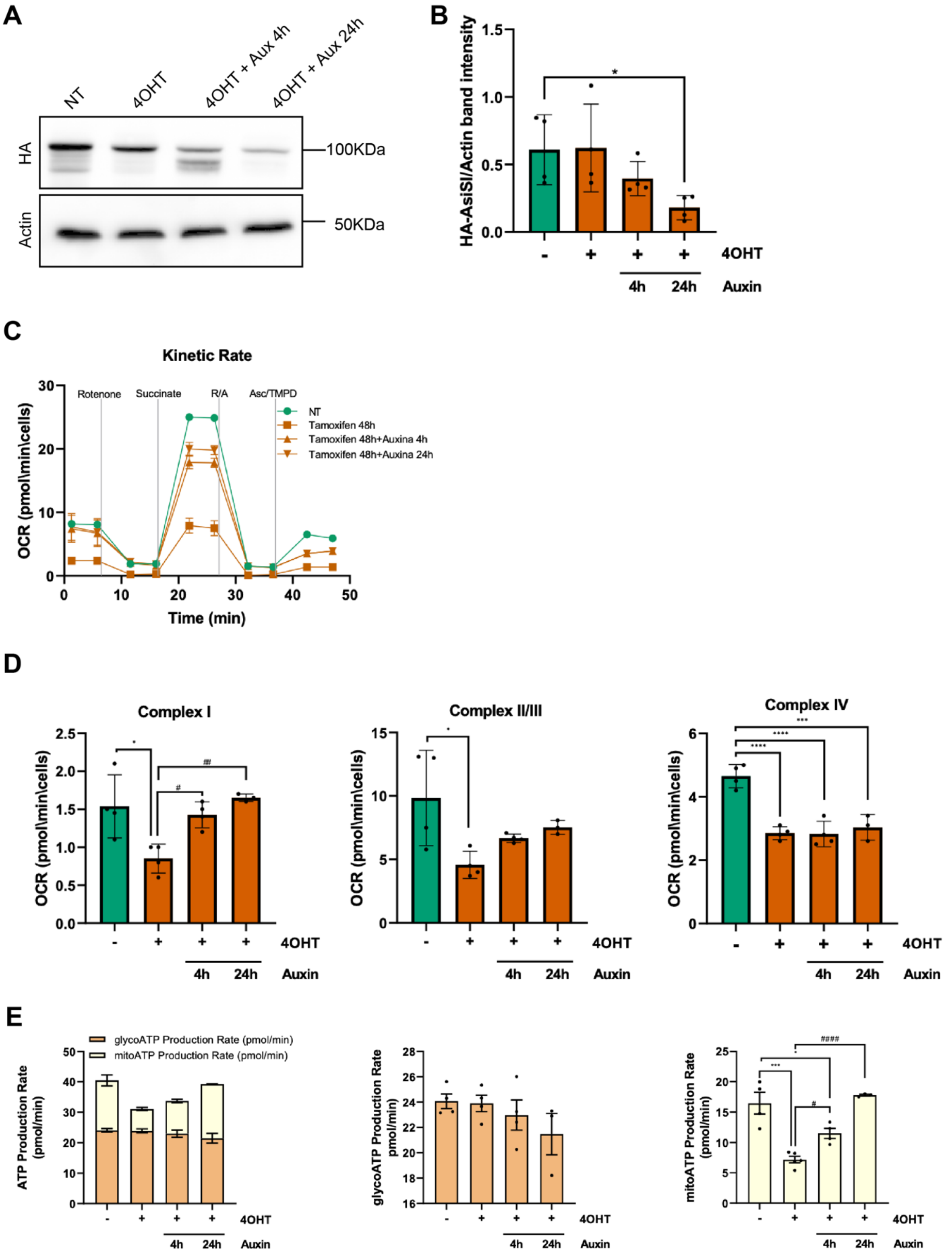


Supplementary Figure 2

**Supplementary Figure 2.**

A. Representative immunoblot showing auxin-induced degradation of the AID-AsiSI-ER fusion protein in AID-DIvA cells following 4-hydroxytamoxifen (4-OHT)-mediated DSB induction and recovery for the indicated times. Actin was used as loading control.
B. Quantification of AID-AsiSI-ER protein levels normalized to Actin following auxin treatment.
C. Representative Seahorse XF Electron Flow Assay profile used to evaluate the activity of individual electron transport chain (ETC) complexes in AID-DIvA cells following DSB induction and recovery upon auxin treatment.
D. Quantification of Complex I, Complex II/III, and Complex IV activities derived from Seahorse XF Electron Flow Assay analyses during DSB recovery.
E. Quantification of mitochondrial and glycolytic ATP production rates obtained by Seahorse XF Real-Time ATP Rate Assay in AID-DIvA cells following DSB induction and recovery after auxin treatment. Data are presented as means ± SEM.

   Statistical significance was determined using one-way ANOVA followed by appropriate post hoc tests. *P < 0.05, ***P < 0.001, ****P < 0.0001 compared with untreated. $^{\#}$P < 0.01, $^{\#\#}$P < 0.001, $^{\#\#\#\#}$P < 0.000.1 compared with 4OHT treated.

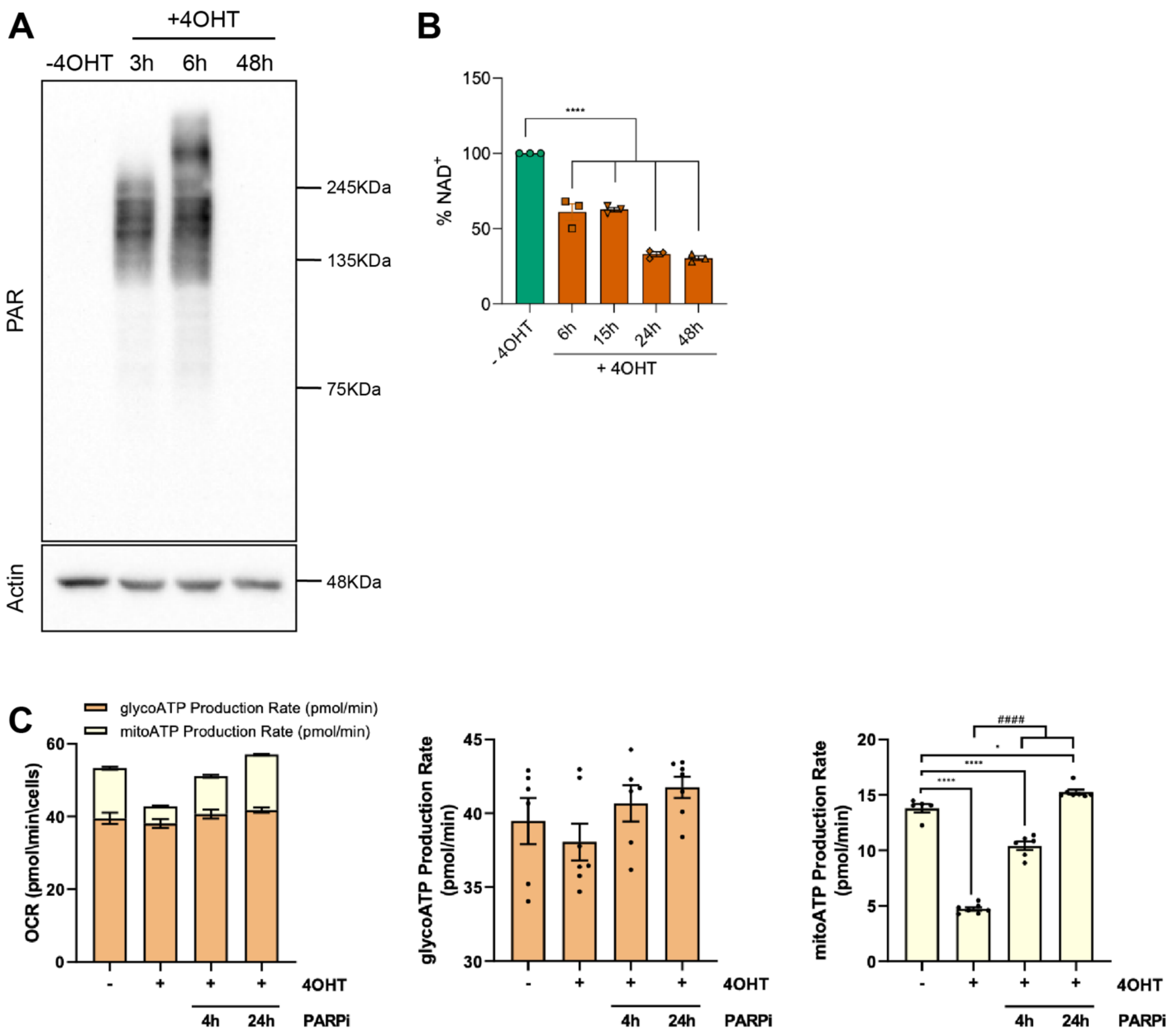


### Supplementary Figure 3

**Supplementary Figure 3.**

A. Representative immunoblot analysis of poly(ADP-ribosyl)ated (PARylated) proteins in AID-DIvA cells treated with 4-hydroxytamoxifen (4-OHT) for the indicated times. Actin was used as loading control.

B. Quantification of intracellular NAD+ levels in AID-DIvA cells following DSB induction by 4-OHT treatment.

C. Quantification of mitochondrial and glycolytic ATP production rates obtained by Seahorse XF Real-Time ATP Rate Assay in untreated and 4-OHT-induced AID-DIvA cells treated with the PARP1 inhibitor PJ34 for the indicated times. Data are presented as means ± SEM.

Statistical significance was determined using one-way ANOVA followed by appropriate post hoc tests. *P < 0.05, ****P < 0.0001 compared with untreated. $^{\#\#\#\#}$P < 0.000.1 compared with 4OHT treated.

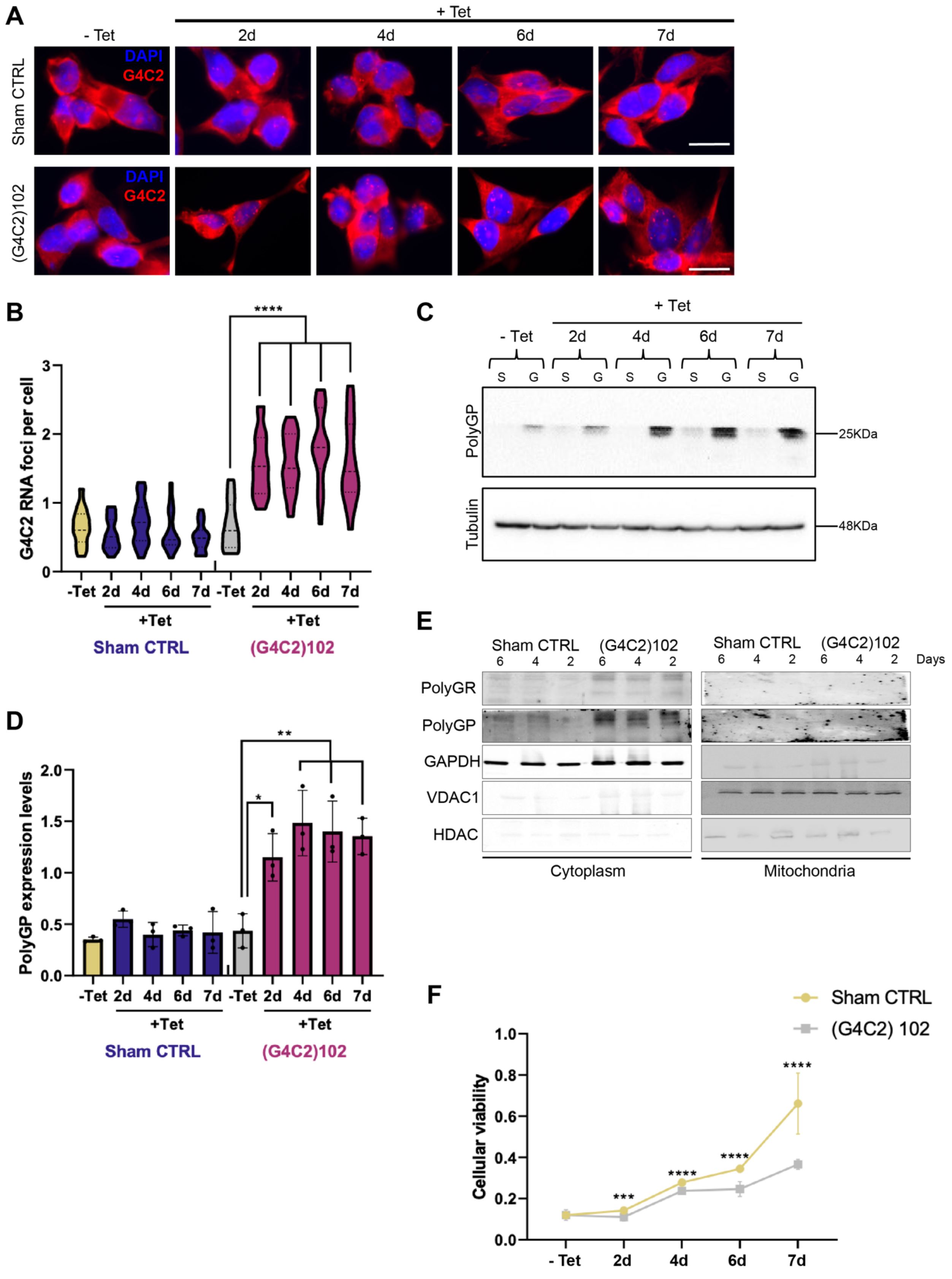


**Supplementary Figure 4**

**Supplementary Figure 4.**

A. Representative fluorescence in situ hybridization (FISH) images showing sense (G4C2) RNA foci in tetracycline-induced NSC-34 sham and NSC-34(G4C2)102 cells. Nuclei were counterstained with DAPI. Scale bar, 20 μm.
B. Quantification of nuclear (G4C2) RNA foci in NSC-34 sham and NSC-34(G4C2)102 cells following tetracycline induction for the indicated times.
C. Representative immunoblot analysis of poly-GP dipeptide repeat (DPR) protein expression in NSC-34 sham control and NSC-34(G4C2)102 cells following tetracycline induction for 2, 4, 6, and 7 days. Tubulin was used as loading control.
D. Quantification of poly-GP DPR expression levels in tetracycline-induced NSC-34 sham and NSC-34(G4C2)102 cells.
E. Representative immunoblot analysis of poly-GR and poly-GP DPR proteins in cytoplasmic and mitochondrial fractions isolated from NSC-34 sham and NSC-34(G4C2)102 cells following tetracycline induction for the indicated times. VDAC1 and HDAC were used as mitochondrial and cytoplasmic fraction markers, respectively.
F. Cell viability analysis of NSC-34 sham and NSC-34(G4C2)102 cells following tetracycline induction for the indicated times. Data are presented as means ± SEM.
   Statistical significance was determined using one-way ANOVA followed by appropriate post hoc tests. *$P < 0.05$, **$P < 0.01$, ***$P < 0.001$, ****$P < 0.0001$.

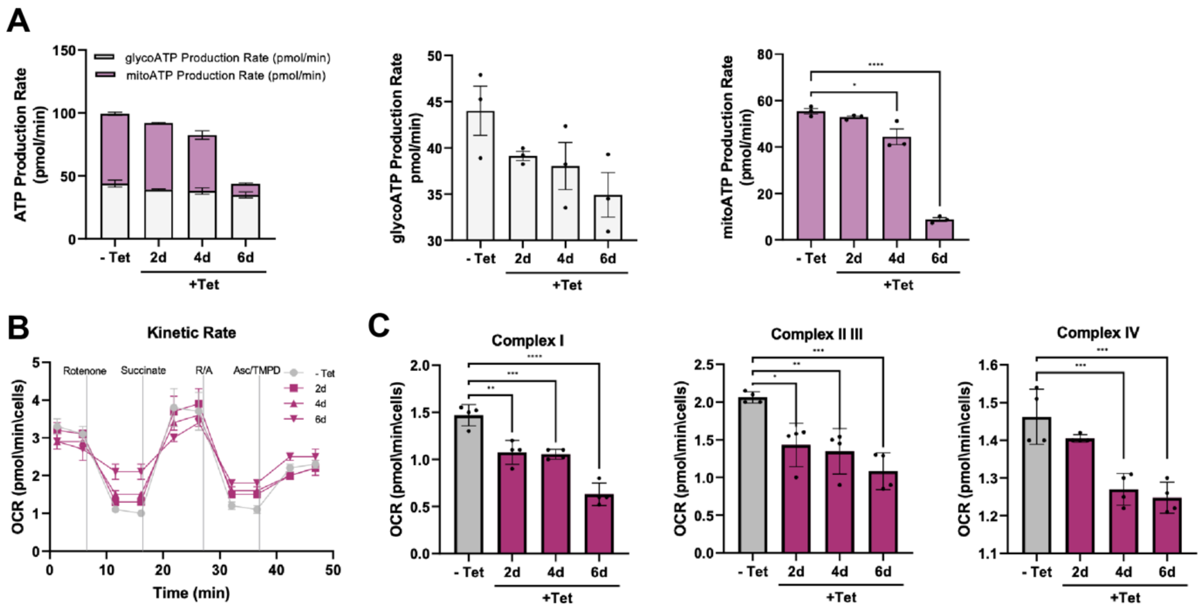


Supplementary Figure 5

**Supplementary Figure 5.**

A. Quantification of mitochondrial and glycolytic ATP production rates obtained by Seahorse XF Real-Time ATP Rate Assay in NSC-34(G4C2)102 cells following tetracycline (Tet) induction for the indicated times.

B. Representative Seahorse XF Electron Flow Assay traces used to evaluate the activity of individual electron transport chain (ETC) complexes in NSC-34(G4C2)102 cells following Tet induction.

C. Quantification of Complex I, Complex II/III, and Complex IV activities derived from Seahorse XF Electron Flow Assay analyses in NSC-34(G4C2)102 cells following Tet induction. Data are presented as means ± SEM.

Statistical significance was determined using one-way ANOVA followed by appropriate post hoc tests. $^*P < 0.05$, $^{**}P < 0.01$, $^{***}P < 0.001$, $^{****}P < 0.0001$.

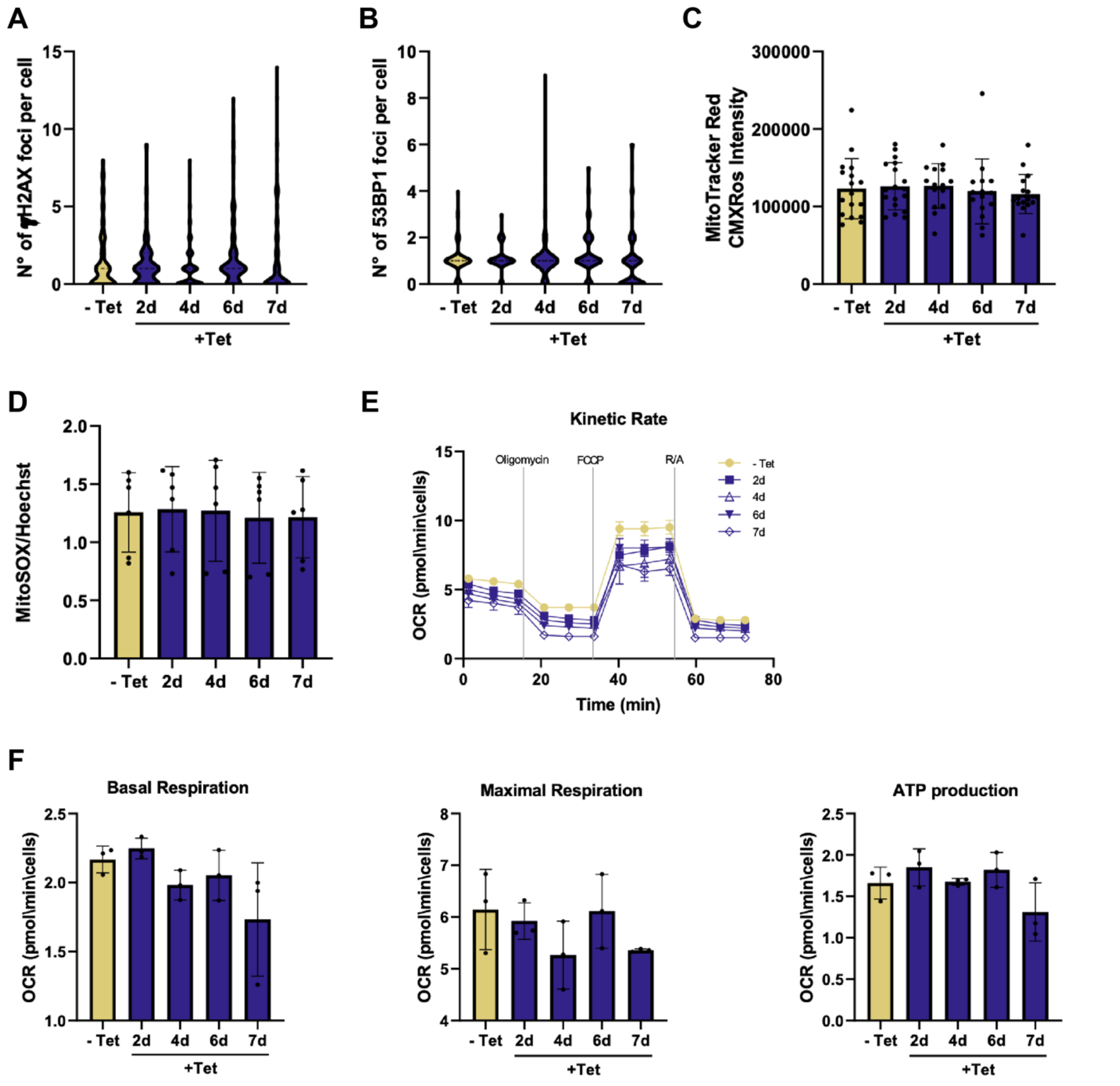


Supplementary Figure 6

**Supplementary Figure 6.**

A. Quantification of γH2AX-positive foci in NSC-34 sham cells following Tet induction.

B. Quantification of 53BP1-positive foci in NSC-34 sham cells following Tet induction.

C. Quantification of MitoTracker Red CMXRos fluorescence intensity in NSC-34 sham cells following Tet induction.

D. Quantification of mitochondrial reactive oxygen species (ROS) production measured by MitoSOX Red fluorescence in NSC-34 sham cells following Tet induction.

E. Quantification of mitochondrial and glycolytic ATP production rates obtained by Seahorse XF Real-Time ATP Rate Assay in NSC-34 sham cells.
F. Additional representative analyses and quantifications related to mitochondrial respiratory function and DDR inhibition in NSC-34 sham cells. Data are presented as means ± SEM. Statistical significance was determined using one-way ANOVA followed by appropriate post hoc tests.

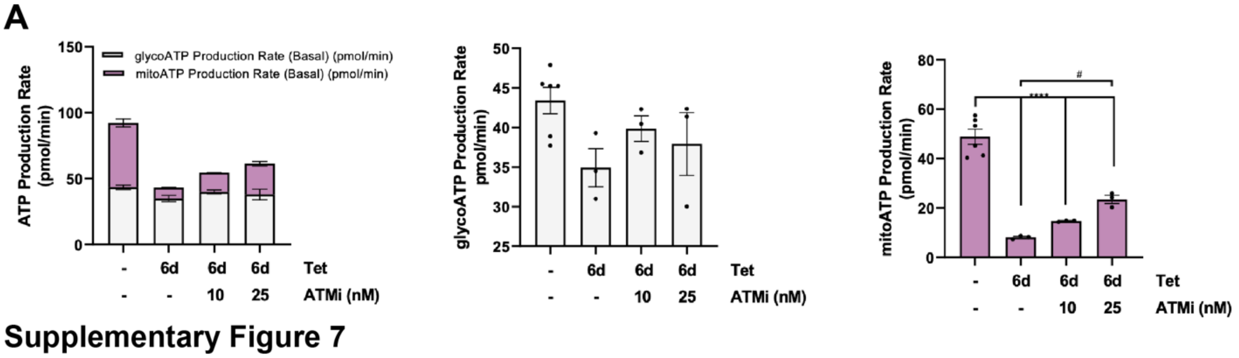


**Supplementary Figure 7.**

A. Quantification of mitochondrial and glycolytic ATP production rates obtained by Seahorse XF Real-Time ATP Rate Assay in NSC-34 (G4C2)102 cells treated with the ATM inhibitor KU-55933 at the indicated concentrations. Data are presented as means ± SEM. Statistical significance was determined using one-way ANOVA followed by appropriate post hoc tests. ****P < 0.0001 compared with untreated. $^{\#}$P < 0.01 compared with 6 days Tet - treated.
